# Gains and losses of the epiphytic lifestyle in epidendroid orchids: review and new analyses with succulence traits

**DOI:** 10.1101/2022.09.30.510324

**Authors:** Géromine Collobert, Benoît Perez-Lamarque, Jean-Yves Dubuisson, Florent Martos

## Abstract

**Background and Aims:** Epiphytism has evolved repeatedly in plants and has resulted in a considerable number of species with original characteristics. Succulent forms in particular are thought to have evolved as an adaptation to the epiphytic environment, because the water supply is generally erratic compared to soils’. However, succulent organs also exist in terrestrial plants, and the question of the concomitant evolution of epiphytism and succulence has received little attention, not even in the epidendroid orchids, which account for 68% of vascular epiphytes.

**Methods:** We reconstructed a new time-calibrated phylogenetic tree of Epidendroideae with 203 genera treated in *Genera Orchidacearum*, from which we reconstructed the evolution of epiphytism and other traits including stem and leaf succulence, while testing the correlated evolution between lifestyle and morphological traits. Furthermore, we reconstructed the ancestral geographic ranges to interpret major character changes during the Cenozoic.

**Key Results:** Epiphytism evolved at least 7.1 My ago in the neotropical Sobralieae, 11.5 My ago in the Arethuseae in Southeast Asia and Australia, and 39.0 My ago in the common ancestor of the Dendrobieae and Cymbidieae in the three previous areas, and was notably lost in the Malaxideae, Collabieae, Calypsoeae, Bletiinae, and Eulophiinae. Stem succulence is inferred to have evolved once, in a terrestrial ancestor 43.1 My ago, thus preceding the evolution of epiphytism by at least 4.1 My. If lost, stem succulence was almost systematically replaced by leaf succulence in epiphytic lineages.

**Conclusions:** Epiphytism probably evolved from terrestrial orchids already possessing succulent stems, which appeared during Eocene climatic cooling. Both epiphytic and secondary terrestrial Epidendroideae may have appeared in seasonally-dry forests. Thus, we believe that the emergence of stem succulence in early epidendroids was a key innovation in the evolution of orchids, facilitating the colonisation of epiphytic environments that led to the greatest diversification of orchids.

## INTRODUCTION

Epiphytism, or the ability of some plants to grow on the surface of other plants, is emerging as a major component of tropical and subtropical forests on Earth (Benzing, 2008; Zotz, 2016). On a global scale, there are currently around 28,000 epiphytic species, *i*.*e*. about 9% of vascular plant diversity (Zotz, 2013, 2016). Epiphytes are located in the Neotropics and in South-East Asia in large numbers, but also in Australia and sub-Saharan Africa (Zotz, 2016; Taylor *et al*., 2022). On a local scale, epiphytes can represent up to 50% of the plant species present in a tropical rainforest (Silvera and Lasso, 2016). Epiphytic diversity ranges across 73 plant families, but three of them concentrate 80% of all vascular epiphytes: Polypodiaceae, Bromeliaceae, and Orchidaceae (Silvera and Lasso, 2016). Orchidaceae account alone for 68% of all epiphytes, conversely 69% of orchid species are epiphytic, and almost all epiphytic orchids belong to one subfamily, the Epidendroideae (Zotz, 2013; Givnish *et al*., 2015).

Despite its ecological importance, epiphytism has been little studied from an evolutionary perspective. We understand not much about the evolutionary patterns of gains and losses of the epiphytic lifestyle within epiphytic lineages, nor the tempo of the physiological or morpho-anatomical traits characterising epiphytic habit, mostly related to capturing and storing water (Silvera and Lasso, 2016; Zotz, 2016; Taylor *et al*., 2022). Indeed, although diverse and heterogeneous in terms of environmental conditions, epiphytic habitats are characterised by an irregular water supply for plants (Benzing, 1987; Zotz, 2016). Adaptation to water shortage in epiphytic habitats could explain why water catching and storing organs are commonly observed among epiphytic plants, such as the tank-forming leaves holding water in some bromeliads (Givnish *et al*., 2014; Zotz, 2016). Most epiphytic orchids and many Araceae have one or more layers of dead epidermal cells surrounding the root, called a velamen, which helps to capture water running off the surface of the tree (Cribb, 1999; Zotz and Winkler, 2013; Stern, 2014). Many epiphytic orchids also have swollen or thickened stems, called pseudobulbs, which store water and nutrients to compensate for irregular supply (Cribb, 1999; Stern, 2014). Leaf succulence is also commonly observed among epiphytes, with or without pseudobulbs in the case of Orchidaceae (Zotz, 2016). In addition, CAM photosynthesis is a water-conserving trait widespread among epiphytic plants (Silvera *et al*., 2009; Zotz, 2016). With approximately 2100 species, the pantropical genus *Bulbophyllum* in Epidendroideae illustrates the ecological and evolutionary success of these characters (Gravendeel *et al*., 2004; Gamisch and Comes, 2019).

However, even though these traits tend to characterise epiphytes, at least some of them are also found in terrestrial plants, with *a priori* no epiphytic ancestors from which they could have retained these characters (Zotz *et al*., 2017). Then, water-deprived traits could have existed in terrestrial ancestors, whose transitions to the epiphytic lifestyle would have been facilitated by the presence of these organs. Rather than adaptations to the epiphytic lifestyle, these traits could have been key innovations (Hunter, 1998; Rabosky, 2017), facilitating the evolution of epiphytism among already drought-adapted terrestrial taxa (in accordance with the assumption of Dressler (1981)). Following Rabosky (2017), we here define a key innovation as ‘the evolution of a trait (or set of functionally linked traits) that provides access to novel resources and that, as a result, facilitates an increase in the total diversification (species richness) of the parent clade’. To note, this definition implies an increase in species richness, but not necessarily in diversification rates.

In a previous study, Givnish *et al*. (2015) investigated the potential drivers of species diversification in the Orchidaceae family. They reconstructed a plastome phylogeny for 75 taxa belonging to 39 species and 16 tribes of orchids, which was then used as a backbone tree for a phylogenetic analysis of 162 species in 18 tribes for which three plastid genes were available. In this way, they were able to demonstrate that several factors, including epiphytic lifestyle and tropical distribution, were associated with higher net diversification rates in orchids. Although they did not specifically address the evolution of epiphytism in the Orchidaceae, their taxon sampling allowed them to identify one transition to the epiphytic lifestyle at the orchid family level, in the largest subfamily Epidendroideae (23,246 species recorded by the World Checklist of Selected Plant Families (WCSP, 2021)). However, they carried this analysis at the level of subtribes only. Two other ancestral estimations of lifestyle evolution were conducted the same year on more extensively sampled phylogenies, including 335 orchid species (Chomicki *et al*., 2015) and 312 Epidendroideae species (Freudenstein and Chase, 2015), respectively. Confirming the early assumption of Dressler (1981), these studies found multiple re-terrestrialization events. Indeed, several lineages nested in the Epidendroideae diversify to the ground and show characters similar to related epiphytic lineages, suggesting that reversions from the epiphytic lifestyle to the terrestrial lifestyle may have occurred in more recent times. Freudenstein and Chase (2015) also detected at least three independent origins of the epiphytism in Epidendroideae, challenging the primary single origin of the epiphytic lifestyle found by Givnish *et al*. (2015) and Chomicki *et al*. (2015). In addition, the ancestral state estimation carried out by Freudenstein and Chase (2015) suggested that the succulent organs common in modern epiphytes could have arisen before the evolution of the epiphytic lifestyle, but the method they used nevertheless returned an uncertain state.

Traditionally, Epidendroideae were divided into basal terrestrial lineages (‘lower Epidendroideae’) and recent epiphytic lineages (‘higher Epidendroideae’). While the latter group includes mainly epiphytic taxa in the tribes Cymbidieae and Vandeae, the former group comprises both terrestrial lineages that are rather basal in the tree and more recent epiphytic lineages, like the Dendrobieae which include the genus *Bulbophyllum*. The studies aforementioned challenge this view. However, extensive uncertainties then remained in the phylogeny of Orchidaceae (Serna-Sánchez *et al*., 2021). In addition, although the phylogeny of Freudenstein and Chase (2015) is to date the best sampled for Epidendroideae, it was not dated, limiting the interpretation of the evolutionary history of the traits. In the light of recent phylogenetic works reappraising the phylogenetic relationships in Epidendroideae using genomic molecular data (Givnish *et al*., 2015; Li *et al*., 2019; Pérez-Escobar *et al*., 2021; Serna-Sánchez *et al*., 2021), there is thus a need to reassess, in a temporal and biogeographical context, the evolutionary history of epiphytism in this subfamily, which contains the vast majority of epiphytic taxa. In addition, new models for ancestral state estimations based on Hidden Markov Models (HMM) have been recently developed and are likely to better reconstruct the evolutionary history of correlated traits (Boyko and Beaulieu, 2021), such as the epiphytic lifestyle and its associated morphological traits.

The purpose of the present study was to analyse in an integrative way the evolutionary history of the epiphytic lifestyle and its associated traits in a single state-of-the-art analysis, using the most recent and robust phylogeny and methods available. (i) We aimed to assess the hypothesis of multiple independent appearances of epiphytism in this orchid subfamily, using a recent backbone tree and the most extended genus sampling possible, while testing for the potential correlated evolution of the epiphytic lifestyle and associated drought-related traits. (ii) We also aimed to ascertain the likelihood of an appearance of succulent traits prior to the epiphytic lifestyle in a climatic and biogeographical framework. (iii) The third objective of this paper was then to characterise the losses of the epiphytic lifestyle in the Epidendroideae and the evolution of the ancestral epiphytic traits.

## MATERIALS AND METHODS

### Phylogenetic analysis of Epidendroideae genera and divergence times

In order to produce a well-sampled, robust, time-calibrated phylogenetic reconstruction of the Epidendroideae, we used a molecular dataset of three plastid genes (*matK, psaB, rbcL*). Sequences were retrieved from GenBank for 203 genera of Epidendroideae listed in *Genera Orchidacearum* (Pridgeon *et al*., 2005, 2009, 2014). Each of the genera was represented in the dataset by sequences obtained from the same voucher specimen (69% of the genera), from different specimens of the same species (14%), or from different species (17%). GenBank accession numbers are provided in Supplementary DataTable S1. Name acceptance by the WCSP (2020) of vouchers and genera was systematically checked. Entirely synonymised genera were merged with the corresponding accepted genus, except for *Cadetia* and *Epigeneium* (synonym: *Dendrobium*) which were kept for the tree calibration (see below). Sequences of each gene were aligned using MAFFT in Geneious Prime 2021.2, manually checked, and then concatenated, resulting in a molecular dataset of 203 Epidendroideae genera and 4649 characters. Best-fitting partition scheme was selected using ModelFinder in IQ-TREE 1.6.12 (Kalyaanamoorthy *et al*., 2017).

A phylogenetic analysis based on these three genes is unlikely to provide strong support for the deep relationships between the genera of Epidendroideae, but tribe relationships in Epidendroideae have been studied several times using 74 to 78 plastid genes (Givnish *et al*., 2015; Li *et al*., 2019; Pérez-Escobar *et al*., 2021; Serna-Sánchez *et al*., 2021). Here we used the multispecies coalescent tree inferred from 78 plastid genes by Pérez-Escobar *et al*. (2021) as a backbone tree for tribe relationships. Genera were assigned to tribes following *Genera Orchidacearum* (Pridgeon *et al*., 2005, 2009, 2014), except for the genera in subtribe Agrostophyllinae (formerly in Vandeae) and *Coelia* (formerly in Epidendreae), following the phylogenetic relationships of Pérez-Escobar *et al*. (2021), and for *Risleya* which was previously misplaced in Malaxideae. Indeed, *Risleya* was found by Xiang *et al*. (2014) to fall within the Collabieae, and by Li *et al*. (2019) within the Calypsoeae. After some tests (data not shown), we found that the phylogenetic position of *Risleya* within Calypsoeae found by Li *et al*. (2019) may be due to undersampling; when the number of sampled taxa was increased, our molecular data supported the position within Collabieae, so we assigned *Risleya* in Collabieae. The backbone tree was input as multiple monophyletic constraints in BEAST 2.6.6 (see below).

The phylogenetic tree was calibrated and inferred in BEAST 2.6.6 (Bouckaert *et al*., 2019), using three fossil calibration points within the Epidendroideae. The stem age of *Earina* was calibrated with the fossil of *Earina fouldenensis*, while the crown age of *Dendrobium* was dated with the fossil of *Dendrobium winikaphyllum* (both dated at 23.2 Mya) (Conran *et al*., 2009). These fossils have the advantage of offering precise calibration points within extant Epidendroideae genera. The crown age of Epidendroideae was calibrated using the fossil of *Succinanthera baltica* estimated at least at 45 Mya (Poinar and Rasmussen, 2017). Despite having only been tentatively assigned to Epidendroideae but not to any extant tribe, this fossil allows to put a minimal age to the Epidendroideae subfamily. Calibration points were set with a log-normal distribution of mean = 1, and of standard deviation (SD) = 1.25 for the two points at 23.2 Mya, and SD = 2 for the crown age of Epidendroideae. The higher SD on the calibration of the root of the Epidendroideae allows the node to take much older values, encompassing the uncertainty of both the age estimation and the phylogenetic position of *Succinanthera baltica*. We set a relaxed uncorrelated log-normal molecular clock, and the birth-death process as tree prior. A diffuse gamma distribution (α = 0.001 and β = 1000) was applied to both clock mean and birth rate priors. Other priors were left as default. Site model averaging using bModelTest 1.2.1 was performed, as recommended by Bouckaert and Drummond (2017), with mutation rate estimated and fixed mean substitution rate. We performed two CoupledMCMC runs of 200 million steps each with eight chains (seven heated, one cold), resampling every 20,000 steps. Stationarity and convergence of runs were assessed using Tracer 1.7.1, and trees were combined using LogCombiner 2.6.6 after discarding the first 12% as burn-in. A maximum clade credibility (MCC) tree with median node heights was calculated using TreeAnnotator 2.6.6.

### Review of drought-related characteristics and lifestyles of the genera of Epidendroideae

To understand the occurrence of the epiphytic lifestyle and of water-capture and water-storage traits in extant Epidendroideae, we reviewed for the 203 extant genera sampled in the phylogenetic analysis the presence of the terrestrial/epiphytic lifestyles. In all, 35 genera also included lithophytic species, but because of the sometimes ambiguous lithophytic lifestyle (Zotz, 2013, 2016), and because only one genus (not sampled in the phylogeny) was scored as entirely lithophytic, we did not include the lithophytic lifestyle, and the genera were then scored as epiphytic and/or terrestrial. We also reviewed the presence of stem succulence (with two modalities if present: heteroblastic (one internode swollen) or homoblastic (several internodes swollen) pseudobulbs/corms), the presence of leaf succulence (including coriaceous leaves), and the minimum and maximum number of velamen layers. Traits were retrieved from *Genera Orchidacearum* (Pridgeon *et al*., 2005, 2009, 2014) and – in particular for the characteristics of the velamen – from *Anatomy of the monocotyledons* (Stern, 2014). Subsequently, we also retrieved data on the shoot architecture of the sampled genera. Indeed, the presence of succulent stems is quite incompatible with the monopodial growth habit (Dressler, 1981). Monopodial shoots grow by unlimited growth of the apical bud, while sympodial shoots have a limited growth and new shoots are produced from an axillary bud (Dressler, 1981). Consequently, the position of the inflorescence is also constrained by the growth form, terminal inflorescences being incompatible with the unlimited monopodial growth, while lateral inflorescences can be encountered in both monopodial and sympodial habits (Dressler, 1981). The position (lateral/terminal) of the inflorescence was then also reviewed.

For all categorical traits, states were mutually non-exclusive, *e*.*g*. genera could include both epiphytic species and terrestrial species. All traits are illustrated in Fig. 1.

**Figure 1.**
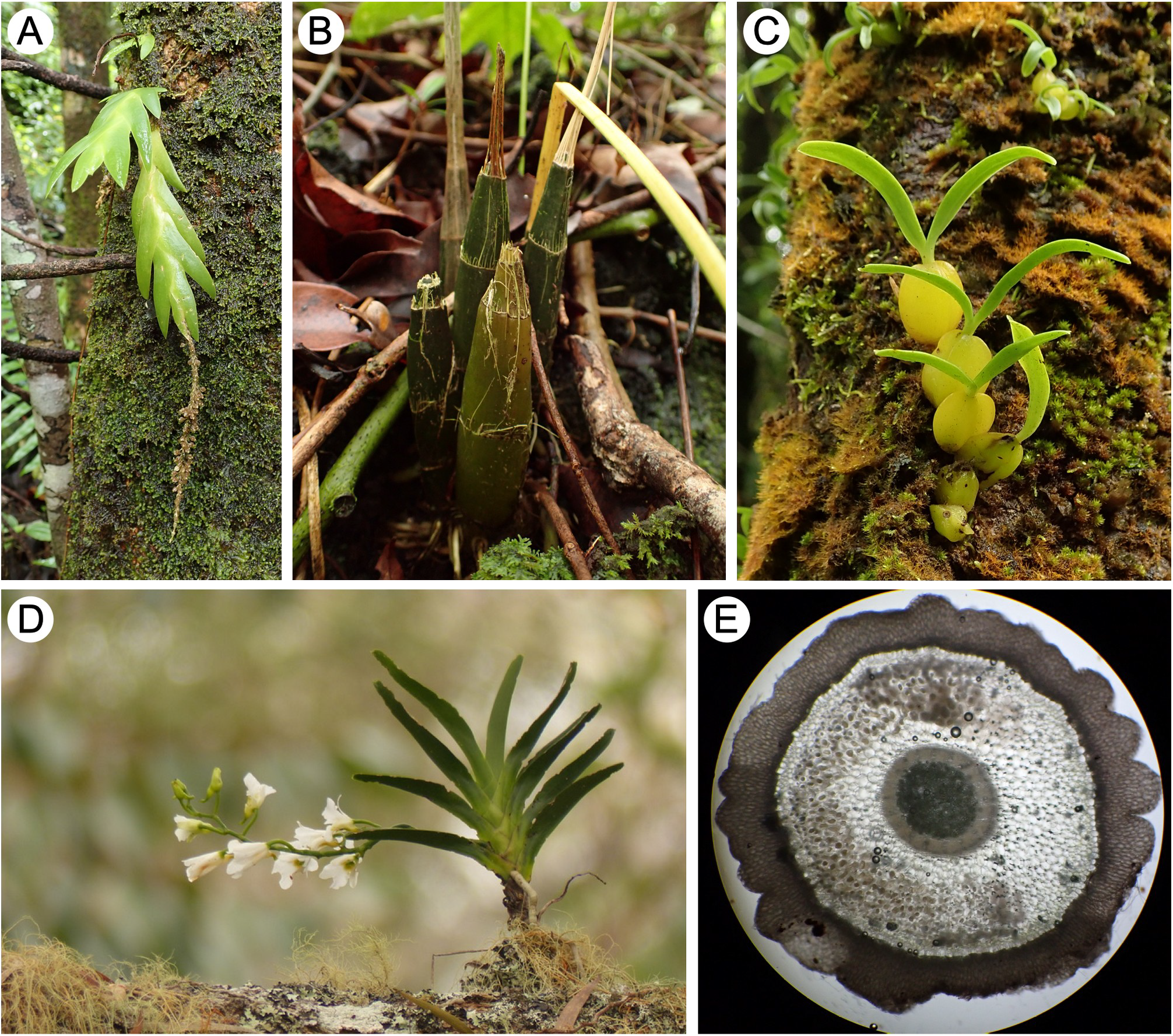
Illustration of morpho-anatomical traits observed among epiphytic and ground-dwelling epidendroid orchids. (A) Succulent leaves, sympodial growth, and terminal inflorescence of the epiphytic *Oberonia disticha* (Malaxideae). (B) Homoblastic pseudobulbs (composed of several internodes) and sympodial growth of the terrestrial *Oeceoclades pulchra* (Cymbidieae; Eulophiinae). (C) Heteroblastic pseudobulbs and sympodial growth of the epiphytic *Bulbophyllum nutans* species complex (Dendrobieae). (D) Succulent leaves, monopodial growth and lateral inflorescence of the epiphytic *Beclardia macrostachya* (Vandeae; Angraecinae), photo courtesy of Rémi Petrolli. (E) Cross-section of a root of *O. pulchra*, showing the brown, multi-layered velamen on the outside of the root.

### Ancestral ranges of the Epidendroideae

Freudenstein and Chase (2015) found three independent origins of the epiphytic lifestyle in Epidendroideae, while Givnish *et al*. (2015) and Chomicki *et al*. (2015) detected only one, but even though Freudenstein and Chase (2015) performed an estimation of the ancestral range (at the Old World/New World level), none of these studies addressed both the time periods and the biogeographic and palaeoclimatic context of the evolution of this lifestyle. Therefore, we estimated the ancestral biogeographic ranges on the time-calibrated MCC tree, using BioGeoBEARS (Matzke, 2013) in R v4.2.1 (R Core Team, 2022). Bioregions were defined as in Givnish *et al*. (2016): North America, Neotropics, Africa, Eurasia, Africa, Southeast Asia, Australia, and Pacific. The models DEC*, DEC*+J, DIVA*, DIVA*+J, BayArea*, and BayArea*+J (excluding a null range) were tested, with a maximal range size of 7, and thus 127 possible states. Time-stratified dispersal multipliers were set following Givnish *et al*. (2016) (Supplementary Data Table S2). The best-fitting model was chosen by AICc comparison and likelihood-ratio tests.

### Test of correlated evolution between epiphytism and succulence traits

The epiphytic habitat is characterised by irregular water-supply. Thus, the epiphytic lifestyle and the succulent organs could have evolved in a correlated way. To test for their potential correlated evolution throughout the phylogeny, we used the discrete dependent/independent approach implemented in BayesTraits v4.0.0 (Pagel and Meade, 2007). For a given trait, we compared the fit of a model where the epiphytic lifestyle and this trait evolved in a correlated way (“dependent model”) to the fit of a model where both traits evolved independently (“independent model”) using Bayes Factors (BF). We ran a reverse-jump (with an exponential prior of mean 10) MCMC analysis for each model, with 5,500,000 iterations, a burn-in of 500,000, and used the stepping stone sampler (Xie *et al*., 2011) with 500 stones and 5000 iterations per stone. Stationarity and convergence of two runs per model were assessed in Tracer v1.7.1. The log BF were computed from the resulting marginal likelihoods, as 2*(log marginal likelihood the dependent model - log marginal likelihood the independent model). We considered a BF > 2 to be significant support for the dependent model.

The test was replicated with another method, i.e. using the R package corHMM v2.7.1, which is a Hidden Markov model designed to allow for the correlated evolution of several characters when estimating transition rates and inferring ancestral states on a phylogeny (Boyko and Beaulieu, 2021). Two symmetric transition matrices were used to correlate or decorrelate transition rates between states (Supplementary Data Table S3). The maximum likelihood of each model was computed with the corHMM function, and best-fitting model was determined by AICc comparison. We considered a AICc difference > 2 between models to be significant support for the model with the lowest AICc. Compared with BayesTraits, corHMM allows some heterogeneity in the transition rates across the whole phylogeny by assuming that the rates can depend on some unobserved (“hidden”) traits.

### Estimation of ancestral lifestyles and drought-related traits

Freudenstein *et al*. (2015) found that succulent stems could potentially have appeared earlier than the epiphytic lifestyle. To ascertain the likelihood of an appearance of succulent traits prior to the epiphytic lifestyle in a temporal framework, transition rates between states were first estimated using maximum likelihood with the corHMM function using 100 random restarts. Polymorphic taxa were included in the analysis, as corHMM can handle multiple states, although this is interpreted as uncertainty by the method rather than polymorphism. We tested models with transition matrices assuming either equal, symmetric, or all-different transition rates, and including or not one hidden rate category. The best-fitting model was determined by AICc comparison. When the AICc difference was lower than 2 between models, the simplest model was preferred over more complex models even if their AICc were slightly lower. Nevertheless, in this case, the more complex models were analysed to verify that they were congruent with the simplest model. Then, 1000 stochastic character maps of the best-fitting model were generated using the makeSimmap function and then summarised to estimate the posterior probability of each state at ancestral nodes. Depending on the results of the test of correlated evolution, morphological traits were also combined with lifestyle in corHMM to take their correlated evolution into account. Transitions between states were estimated from the posterior probability distribution of trait changes from the stochastic character mapping.

For the mean number of velamen layers in the root, we considered the trait to be continuous. We used the function anc.ML from the R package phytools (Revell, 2012) with a Brownian Motion model of continuous character evolution to estimate the ancestral states. The pgls function of the R package caper (Orme *et al*., 2018) was used to test if the mean number of velamen layers in epiphytic taxa is significantly different from that of terrestrial taxa, while accounting for the phylogenetic relatedness between species. As several genera are both epiphytic and terrestrial, we repeated the PGLS ten times, each time randomly assigning each polymorphic genus to be only epiphytic or terrestrial.

## RESULTS

### Phylogenetic relationships and divergence times between Epidendroideae genera

All epidendroid subtribes *sensu* Genera Orchidacearum were found monophyletic in the obtained phylogenetic tree of 203 genera of Epidendroideae, excepting the subtribe Cymbidinae which was split into three clades (Fig. 2). According to our time-calibration based on three fossils (including a minimal age for the Epidendroideae based on the fossil described in Poinar and Rasmussen (2017)), the crown age of Epidendroideae was estimated at 47.4 Mya (95% Highest Posterior Density (95% HPD) interval = 63.2–45.0 Mya). Tribe crown ages and corresponding 95% HPD intervals are summed up in Table 1.

**Figure 2.**
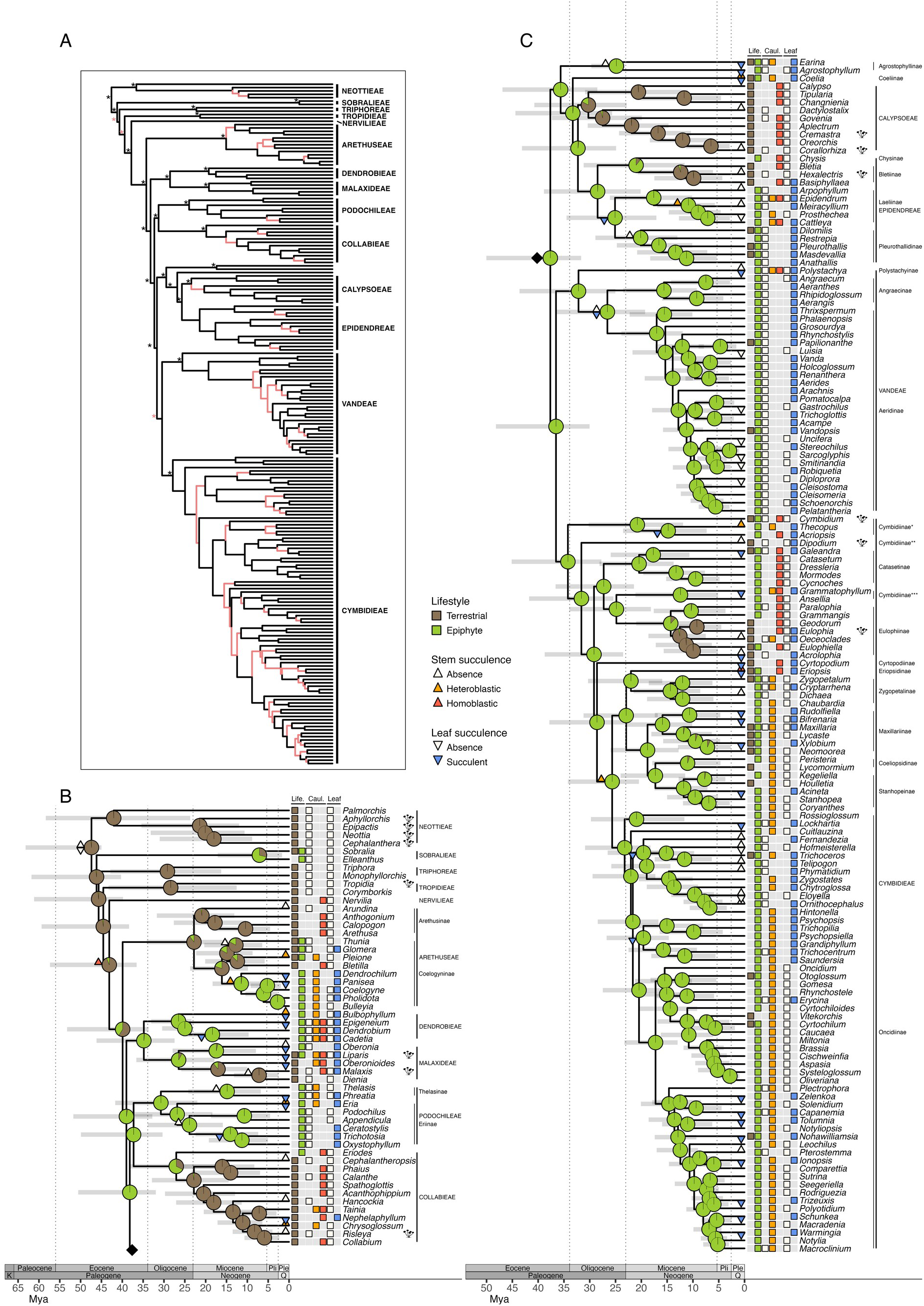
Evolution of lifestyle and succulent forms (stem and leaves) in epidendroid orchids. (A) Dated phylogenetic tree reconstructed from 203 genera in 14 tribes of epidendroid orchids. Branches with posterior probability < 0.95 are coloured coral-red. Nodes that were constrained in the analysis are indicated by stars (in red when the quartet support for the main topology was < 50% in the coalescent-species tree of Pérez-Escobar *et al*. 2021). (B) and (C) Estimation of ancestral states in the Epidendroideae genus tree with corHMM. As lifestyle and leaf succulence were found to be correlated (see these results in Table 2), these traits were estimated at once using a correlated transition matrix in corHMM. Pie charts at nodes represent ancestral lifestyles and their probabilities. For succulence traits, only significant state changes, i.e. state probability > 0.5 while < 0.5 in the ancestral node, were indicated by coloured triangles (see Supplementary Data Figs. S6 and S8 for full estimations of the ancestral states of stem and leaf succulence respectively). The character states of the present genera are represented on the right side of the tree by coloured boxes. In addition, genera with mycoheterotrophic species are indicated by a mushroom symbol. Cymbidiinae are split into three clades which are indicated by one, two and three stars after the subtribe name.

**Table 1.** Estimated crown age and 95% highest posterior density (95% HPD) interval of epidendroid orchid tribes from our study *versus* Givnish *et al*. (2015) and Serna-Sánchez *et al*. (2021). Values from Serna-Sánchez *et al*. (2021) were retrieved from their relaxed molecular clock, birth-death divergence time analysis. Empty cells are tribes for which only one genus was sampled and therefore crown ages are not available. As Nervilieae are represented by only one genus in the three studies, their crown age is not available and therefore they are not included in the table. Our crown age estimates are within the 95% HPD intervals found by Givnish *et al*. (2015) and Serna-Sánchez *et al*. (2021) for all orchid tribes, except for Neottieae, for which our age estimate is significantly younger than the 95% HPD interval in Serna-Sánchez *et al*. (2021).

| Orchid tribe | Crown age [95% HPD]<br>(Mya) in this study | Crown age [95% HPD]<br>(Mya) in Givnish *et al.*<br>(2015) | Crown age [95% HPD]<br>(Mya) in Serna-Sánchez *et al.* (2021) |
| --- | --- | --- | --- |
| Arethuseae | 22.8 [34.1–14.7] | 19.1 [25.2–13.6] | 21.7 [34.8–12.6] |
| Calypsoeae | 30.3 [40.7–22.8] | 30.8 [35.1–26.7] | 33.1 [40.4–26.1] |
| Collabieae | 27.1 [38.2–16.8] | 23.2 [29.7–16.1] | 22.1 [35.1–10.7] |
| Cymbidieae | 34.2 [45.1–27.7] | 30.7 [34.9–26.5] | 35.3 [42.9–28.2] |
| Dendrobieae | 26.5 [33.1–23.4] | 22.0 [29.9–13.4] | 24.3 [29.5–19.4] |
| Epidendreae | 28.5 [38.9–20.1] | 25.5 [31.5–18.4] | 31.9 [38.9–25.0] |
| Malaxideae | 26.5 [36.6–17.9] | 23.5 [30.1–17.7] | 28.0 [37.1–18.0] |
| Neottieae | 42.1 [58.4–23.7] | 44.6 [52.2–37.1] | 56.5 [**69.2–43.1**] |
| Podochileae | 30.7 [41.7–22.0] | 27.9 [33.3–21.6] |  |
| Sobralieae | 7.1 [17.1–1.7] | 10.3 [20.8–3.7] | 10.9 [19.0–5.7] |
| Triphoreae | 29.2 [42.9–16.3] | 28.0 [37.2–19.0] |  |
| Tropidieae | 28.4 [43.3–12.6] | 34.9 [44.0–25.0] |  |
| Vandaeae | 32.2 [43.6–25.0] | 27.3 [32.4–22.0] | 30.8 [39.3–22.2] |

### Lifestyles and drought-related traits among genera of Epidendroideae

We compiled data for the 203 genera sampled in the phylogeny, of which 201 were accepted and 2 were synonyms according to the WCSP (2020). Among the 201 accepted genera, 126 (62.7%) were epiphytic, 50 (24.9%) were terrestrial, and 25 (12.4%) were both epiphytic and terrestrial. Stem succulence was found in 104 genera (51.7%), while 22 genera (10.9%) were polymorphic. Leaf succulence was present in 74 genera (36.8%), while 21 genera (10.4%) were polymorphic. Among the 126 genera (62.7%) lacking succulent leaves, 14 were mycoheterotrophic and thus leafless. Statistics for all traits are summep up in Supplementary Data Table S4. Traits of the velamen were missing for 66 genera out of 201 in the phylogeny (32.8%).

### Ancestral ranges of Epidendroideae

The best-fitting model was a BayAreaLike* model with jump speciation (Supplementary Data Table S5), and the difference with the BayAreaLike* model without jump speciation was significant according to the likelihood-ratio test (Supplementary Data Table S6). The most recent common ancestor (MRCA) of Epidendroideae was inferred to range from the Neotropics to Southeast Asia and Australia (Fig. 3). In Neottieae and Calypsoeae, taxa dispersed northwards up to Eurasia and North America during the Miocene. Triphoreae, Epidendreae, the subtribe Catasetinae and all genera from *Cyrtopodium* to *Notylia* in Cymbidieae mostly withdrew to the Neotropics during the Oligocene, as well as Sobralieae during the late Miocene. In Epidendreae, the MRCA of Bletiinae spread to North America 12.4 My ago (probability of state p = 0.91, node posterior probability PP = 1, 95% HPD = 20.7–6.1 Mya). Nervilieae, Arethuseae, Podochileae, Collabieae, Agrostophyllinae, and Aeridinae withdrew independently mostly to South Asia and Australia during the Oligocene and the Miocene. In Dendrobieae and Malaxideae, genera mostly spread northwards to Eurasia, as well as to Africa and to the Pacific islands. Finally, the subtribes Angraecinae and Eulophiinae diversified in Africa during the Miocene.

**Figure 3.**
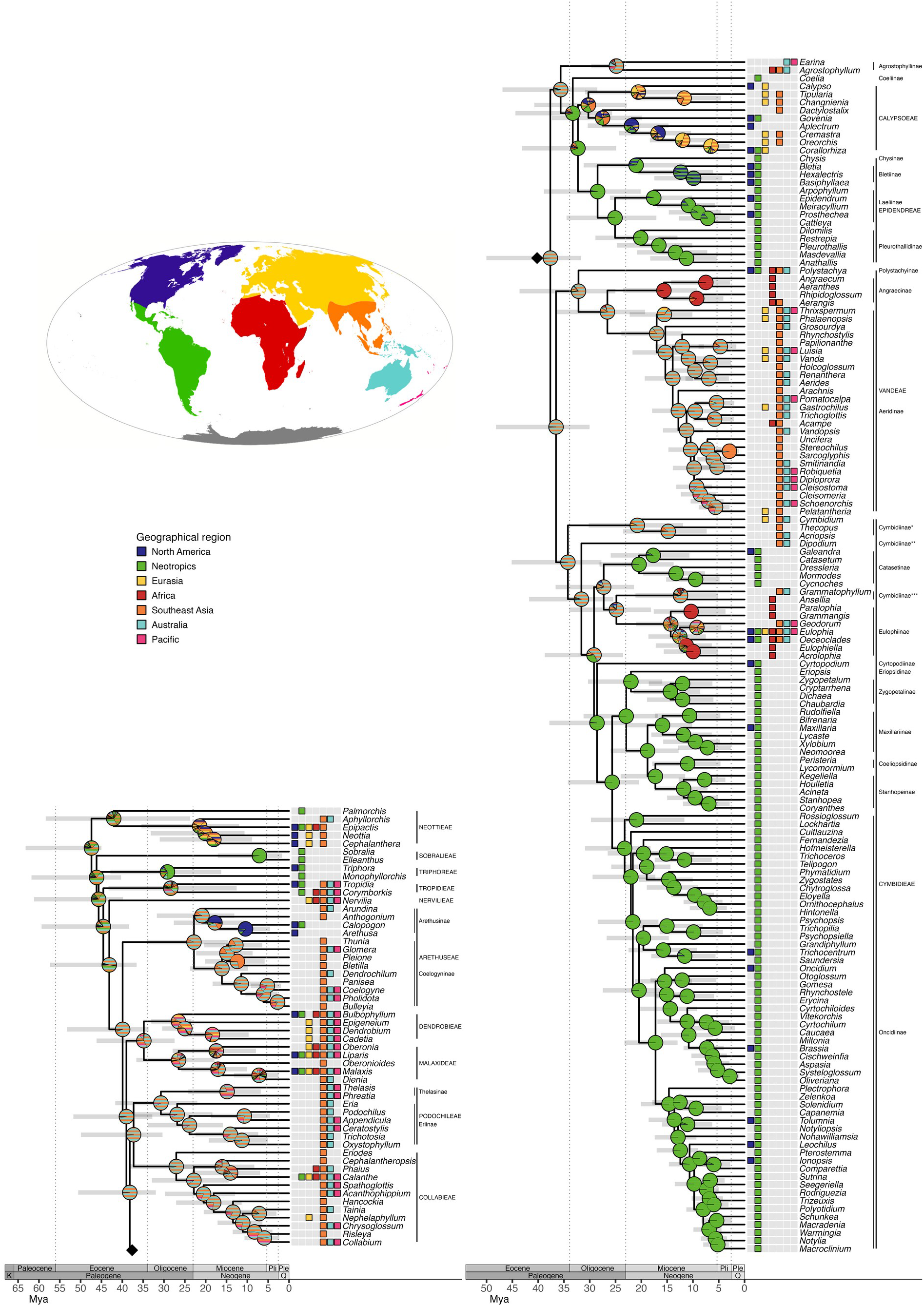
Estimation of ancestral geographic ranges in the dated phylogenetic tree of the 203 genera of epidendroid orchids, using the BayAreaLike*+J model in BioGeoBEARS. The different geographical areas occupied by the present-day Epidendroideae genera are represented by unique colours. The pie charts at nodes represent the ancestral geographic ranges in proportion to their likelihood. These are solid-coloured when the range comprises a single area (*e*.*g*. Neotropics), or striped when the range comprises several geographical areas (*e*.*g*., a range represented by orange and light blue bands includes both Southeast Asia and Australia). The geographical ranges of extant genera are represented on the right-hand side of the tree by coloured boxes.

### Test of correlated evolution between epiphytism and succulence traits

Log marginal likelihoods of dependent/independent models computed with BayesTraits, as well as the resulting log BF, are shown in Table 2. All morphological traits were found to be correlated to the lifestyle. Very strong evidence of correlated evolution was found between the lifestyle and leaf succulence, and between the lifestyle and growth form. Only these combinations showed a difference of AICc > 2 for correlated evolution with corHMM models (Table 2). Therefore, because leaf succulence and growth form were found to be evolutionary correlated with the lifestyle by both methods, we thus jointly estimated their evolutionary histories with the lifestyle (using correlated matrices of transition between states) in corHMM. Conversely, because the stem succulence and the position of inflorescence were not found to be evolutionary correlated with the lifestyle according to corHMM, we estimated alone their evolutionary histories using corHMM (see below).

**Table 2.** Tests of correlated evolution between orchid genus life style (epiphytic/terrestrial) and stem succulence (absence/presence), leaf succulence (absence/presence), inflorescence position (lateral/terminal), and growth form (monopodial/sympodial), using two methods (BayesTraits and corHMM). The log marginal likelihood (LML) of the dependent or independent evolution models, and the log Bayes factor (BF), were calculated using BayesTraits. Symmetric matrices (SYM) were used in corHMM to correlate or decorrelate the two traits (see Supplementary Data Table S3 for more details on these matrices). A delta AICc between 0 and 2 indicates that the AICc values are not significantly different, in which case the simpler (uncorrelated) model is preferred. A delta AICc > 2 indicates that the correlated model fits the data significantly better. A delta AICc < -2 indicates that the uncorrelated model fits the data significantly better.

|  | BayesTraits |  |  |  | corHMM |  |  |
| --- | --- | --- | --- | --- | --- | --- | --- |
| Tested correlation | LML of independent model | LML of dependent model | log BF | Interpretation | AICc correlated SYM matrix | AICc uncor-related SYM matrix | delta AICc |
| Lifestyle~Stem succulence | -147.4 | -144.0 | 6.7 | Strong evidence | 276.9 | 278.8 | 1.9 |
| Lifestyle~Leaf succulence | -183.1 | -172.8 | 20.7 | Very strong evidence | 324.4 | 344.5 | 20.2 |
| Lifestyle~Inflorescence | -131.5 | -127.5 | 8.1 | Strong evidence | 246.9 | 244.6 | -2.4 |
| Lifestyle~Growth form | -104.3 | -98.7 | 11.2 | Very strong evidence | 185.8 | 194.5 | 8.7 |

### Estimation of ancestral lifestyles (terrestrial or epiphytic)

The evolutionary history of the epiphytic lifestyle was very consistent across models, i.e. either with the uncorrelated model (Supplementary Data Fig. S1) or with models correlated to leaf succulence (Fig. 2) or to growth form. These models inferred that the MRCA of Epidendroideae was terrestrial (p = 0.99). We found that the epiphytic lifestyle likely appeared at least three times independently in ancestral nodes, namely in (i) the MRCA of Dendrobieae and Cymbidieae 39.0 My ago (p = 0.98, node PP = 1 (backbone node), 95% HPD = 51.7–32.8 Mya), (ii) in the MRCA of *Dendrochilum* and *Panisea* in Arethuseae 11.5 My ago (p = 0.98, node PP = 0.98, 95% HPD = 18.3–6.3 Mya), and (iii) in the MRCA of Sobralieae 7.1 My ago (p = 0.71, node PP = 1 (backbone node), 95% HPD = 17.1–1.7 Mya).

Likewise, we inferred at least five transitions from the epiphytic lifestyle to terrestriality (secondary terrestrialization) leading to diversification in (i) the MRCA of *Oberonioides* and *Malaxis* in Malaxideae 17.0 My ago (p = 0.89, node PP = 1, 95% HPD = 25.9–10.1 Mya), (ii) in the Collabieae excluding *Eriodes* 22.8 My ago (p = 0.97, node PP = 1, 95% HPD = 32.9–14.5 Mya), (iii) in the MRCA of Calypsoeae 30.3 My ago (p = 0.83, 95% HPD = 40.7–22.8 Mya), (iv) in the MRCA of Bletiinae 12.4 My ago (p = 0.95, node PP = 1, 95% HPD = 20.7–6.1 Mya), and (v) in the MRCA of *Eulophia* and *Oeceoclades* 12.6 My ago (p = 0.97, node PP = 0.83, 95% HPD = 18.6–7.5 Mya).

Using the posterior probability distribution of trait changes from stochastic mapping of the correlated model of lifestyle and growth form, transitions from the terrestrial to the epiphytic lifestyle were estimated to have occurred three to 11 times (95% HPD interval) in the phylogeny with a median of five changes. Reverse transitions i.e., from the epiphytic to the terrestrial lifestyle, were estimated to have occurred eight to 17 times (95% HPD interval) in the phylogeny, with a median of 13.

Our analysis revealed that the lifestyle constrained the growth form in Epidendroideae. Indeed, no genus in the phylogeny was scored as terrestrial and monopodial without being polymorphic for at least one trait. Thus, except for *Maxillaria* which was polymorphic for both the lifestyle and the growth form, all transitions from the ancestral sympodial growth (p = 1) towards the monopodial growth occurred in entirely epiphytic taxa (Fig. 2, Supplementary Data Fig. S2). Consequently, transition rates are close to zero between monopodial epiphytes and monopodial terrestrials and between sympodial terrestrials and monopodial terrestrials (Supplementary Data Fig. S3). Transitions between terrestrial and epiphytic lifestyles occurred within sympodial taxa, at a rate of 0.0064 event/My, and afterward transitions to a monopodial growth occurred within epiphytic taxa at a rate of 0.0039 event/My (Supplementary Data Fig. S3). Using the posterior probability distribution of trait changes from stochastic mapping of the correlated model of lifestyle and growth form, throughout the phylogeny transitions to monopodial growth occurred 12 times at the most (95% HPD [6, 10], median = 8), with only one transition leading to diversification (in Vandeae) and 11 transitions at tips, of which ten occurred in Cymbidieae (Supplementary Data Fig. S2). As the position of the inflorescence is constrained by the growth form, as expected we found that all monopodial taxa evolved lateral inflorescences (Supplementary Data Fig. S4), but no correlation was undoubtedly found with the evolution of the epiphytic lifestyle (Table 2). Full description of the evolution of the position of inflorescence is available in Supplementary Data.

### Estimation of ancestral drought-related traits (stem succulence, leaf succulence and velamen)

Ancestral state estimation of stem succulence was consistent either when considering the presence-absence of succulence (Supplementary Data Fig. S5) or when dividing the presence of stem succulence into heteroblastic (one internode swollen) or homoblastic (several internodes in succession swollen) (Fig 2, Supplementary Data Fig. S6). The absence of stem succulence was inferred as ancestral in Epidendroideae (p = 0.80). Stem succulence was inferred to be ancestrally homoblastic, and to have appeared only once, in the MRCA of Nervilieae and Cymbidieae 43.1 My ago (p = 0.82, node PP = 1 (backbone node), 95% HPD = 57.3–36.6 Mya) (Fig 2, Supplementary Data Fig. S6), thus predating the epiphytic lifestyle by at least 4.1 My. The heteroblastic stem succulence evolved repeatedly from homoblastic pseudobulbs at least eight times throughout the phylogeny (distribution of changes from stochastic mapping: 95% HPD [9, 16], median = 12) (Supplementary Data Fig. S6). Stem succulence appeared to have been lost several times, notably in the MRCA of Vandeae excluding *Polystachya* 26.6 My ago (p = 0.86, node PP = 1, 95% HPD = 36.5–19.3 Mya), when simultaneously growth form changed from sympodial to monopodial (p = 0.946) (Supplementary Data Fig. S2). Transition rates (Supplementary Data Fig. S7) indicated that the stem succulence was fifteen-fold more often lost (0.033 event/My) than acquired (0.0022 event/My), and confirmed that homoblastic pseudobulbs tended to be either lost (0.02 event/My) or to evolve into heteroblastic pseudobulbs (0.011 event/My). Heteroblastic stem succulence derived predominantly from homoblasty and was lost as often as it was acquired (0.013 and 0.011 event/My, respectively), though mostly at tips (Supplementary Data Fig. S6).

Leaf succulence (Supplementary Data Fig. S8) was inferred to have appeared several times convergently (distribution of changes from stochastic mapping: 95% HPD [33, 52], median = 43 in epiphytic taxa; 95% HPD [77, 112], median = 93 in terrestrial taxa), mostly at tips, and particularly during Oligocene in epidendroid lineages that were probably already epiphytic at that time (*i*.*e*. in Podochileae, Epidendreae, Vandeae, and at least twice in Cymbidieae) (Fig. 2, Supplementary Data Fig. S8). However, even though leaf succulence evolved predominantly in epiphytic groups, some terrestrial taxa also evolved it, for example in genera *Oberonioides, Acrolophia*, or *Cyrtopodium* (Fig. 2, Supplementary Data Fig. S8). In Vandeae, the appearance of leaf succulence 26.6 My ago (p = 0.76, node PP = 1, 95% HPD = 36.5–19.3 Mya) was inferred to coincide with the loss of stem succulence and the change from sympodial to monopodial growth (Fig. 2, Supplementary Data Fig. S2).

The number of velamen layers (Supplementary Data Fig. S9) tended to be ancestrally low in Epidendroideae, and seems to have increased in the MRCA of *Panisea* and *Bulleyia* in Arethuseae, in Dendrobieae, in Collabieae, in Laeliinae, and in most of Cymbidieae, as well as in *Govenia* (tribe Calypsoeae). In Vandeae, most genera miss data on velamen, thus inheriting ancestral state without change, hence some genera may actually have a high number of velamen layers in this tribe. As expected, primarily terrestrial tribes tended to have only a few velamen layers, but some secondary terrestrial genera, *i*.*e. Govenia, Cyrtopodium*, and the subtribe Eulophiinae have among the highest numbers of layers. All PGLS models indicated that terrestrial taxa tended to have fewer velamen layers than epiphytic ones, but a high number of velamen layers was not significantly associated with epiphytic lifestyle (range of F-statistic [2.2–0.099] on 1 and 135 DF, p-value [0.14–0.75]).

## DISCUSSION

### A consistent time-calibrated, phylogenetic and biogeographical framework

Our sampling increases by 110 genera the sampling in Givnish *et al*. (2015) and by 111 the sampling of Serna-Sánchez *et al*. (2021) and Pérez-Escobar *et al*. (2021). The root age of Epidendroideae was similar to the age found by Givnish *et al*. (2015) (48.05 Mya, 95% HPD = 55.7–40.7 Mya), even if the HPD interval is left-shifted in our analysis, due to the log-normal calibration parameters. All the crown ages of the tribes were comprised in the 95% HPD interval obtained by Givnish *et al*. (2015), but not in all the intervals obtained from the relaxed birth-death tree of Serna-Sánchez *et al*. (2021) (Table 1). Subtribe relationships in Cymbidieae were congruent with the plastid ML tree of Serna-Sánchez *et al*. (2021) and of Pérez-Escobar *et al*. (2021), except for the position of the Cyrtopodiinae, which were placed sister to the Catasetiinae with high support in their studies, whereas in our study the position of the Cyrtopodiinae is not supported (node PP = 0.34, see Fig. 2). Subtribe relationships seem more unresolved in the Epidendreae, with incongruences between all studies. Our subtribe relationships in this tribe were congruent with Serna-Sánchez *et al*. (2021), but not with Pérez-Escobar *et al*. (2021), however node supports were lower in this last study.

The results of our estimation of ancestral biogeographic ranges in the Epidendroideae were congruent with those of Givnish *et al*. (2016), although the most recent common ancestor (MRCA) of Epidendroideae was inferred to range from the Neotropics to Southeast Asia and Australia, when Givnish *et al*. (2016) found it to range only in the Neotropics or in the Neotropics and Southeast Asia at the same time. The restriction of the range of taxa to either Southeast Asia and Australia, or to the Neotropics, in almost all tribes independently during the Oligocene (Fig. 3), and the build-up of polar ice over Antarctica up to the glaciation event at the Eocene-Oligocene boundary seems to support the hypothesis that the Neotropics and the Australasian region could have previously been connected via Antarctica (Bush *et al*., 2011). The ice barrier probably led to allopatric speciation as it could have been the case in Cymbidieae and maybe also between Calypsoeae and Epidendreae, or to extinction in one of these regions.

### Where and when did the epiphytic lifestyle of Epidendroideae evolve?

In agreement with Freudenstein and Chase (2015), we found that the epiphytic lifestyle likely appeared at least three times independently in ancestral nodes, against one single origin in Givnish *et al*. (2015) and Chomicki *et al*. (2015). The first occurrence of the epiphytic lifestyle was inferred at the end of the Eocene in a range encompassing the Neotropics, Southeast Asia, and Australia (Figs. 1 and 2). It is likely that epiphytism arose in seasonal forests, maybe primarily on cliffs and rocky areas, as hypothesised by Dressler (1981).

Indeed, the major retractation of rainforests under the cooling climate during the Eocene (Bush *et al*., 2011) could have directed the evolution of epiphytism in seasonal forest canopies. However, contrary to Dressler (1981) who supposed that species adapted to the epiphytic habitats by developing succulent organs, we rather conclude that epiphytism evolved among already drought-adapted taxa, and thus the ancestral stem succulence (Fig. 2) may have been an important feature for the evolution of epiphytism (see next section). Indeed, Frenzke *et al*. (2016) also hypothesised that the terrestrial ancestor of epiphytic *Peperomia* already showed traits potentially facilitating epiphytic life such as succulence or CAM metabolism.

In Arethuseae, epiphytism likely appeared during the Miocene in the Southeast Asian and Australian range, when the climate was cooling again after the Mid-Miocene climatic optimum and moist megathermal forests were restricted to the tropical zone, with the exception of the Australasian region (Bush *et al*., 2011). In Southeast Asia, everwet climates predominated, but some regions could have been wet but seasonal, with open vegetation (Bush *et al*., 2011). Thus, it is unclear if epiphytic Arethuseae most likely appeared in seasonal or everwet forests, and in addition the ancestral stem succulence could have facilitated the transition to the epiphytic habitat in both environments. Moreover, Coelogyninae are at present predominantly inhabiting everwet forests, and occur less frequently in areas with seasonal climates (Pridgeon *et al*., 2005).

Interestingly, our findings that the epiphytic lifestyle likely evolved from drought-adapted plants contrast with the inferred evolution of other epiphytic lineages, which were rather found to have diversified mostly in rainforests or moist montane forests (Wikström *et al*., 1999; Schuettpelz and Pryer, 2009; Calvente *et al*., 2011; Givnish *et al*., 2014; Bechteler *et al*., 2021; Chen *et al*., 2022). This could be explained by the absence of such pre-existing drought-tolerant traits in these groups. Indeed, even among the Epidendroideae, the last appearance of the epiphytic lifestyle likely occurred in Sobralieae in absence of any type of succulence 7.1 My ago, at the end of the Miocene, in the Neotropics. In Bromeliaceae, subfamily Bromelioideae (tank bromeliads), epiphytism evolved around 5.9 Mya in the late Miocene, in the Atlantic forest region of Neotropics, during a global cooling of the climate, synchronously with the uplift of the Serra do Mar which would have favoured cooler, rainier, more humid conditions in the Atlantic forest region (Givnish *et al*., 2014). Moreover, humid montane habitats were found to have favoured both epiphytic evolutions in Bromeliaceae (Givnish *et al*., 2014). Thus, considering the several mountain uplifts in the Neotropics since the Miocene (Potter and Szatmari, 2009; Givnish *et al*., 2014; Martins *et al*., 2018), and that Sobralieae lack succulent organs, epiphytism in this tribe may also have evolved in humid montane habitats.

Nevertheless, the hypothesis of an evolution of epiphytism in everwet forests for all occurrences of the epiphytic lifestyle should not be rejected: in Epidendroideae the pre-adaptation to drought would also have been beneficial in everwet habitats. Indeed, CAM photosynthesis, a water-conserving metabolism, could have been selected in high rainfall habitats in *Bulbophyllum* (Gamisch *et al*., 2021), and many species of Epidendroideae with succulent organs now grow in the shade of everwet forests (Pridgeon *et al*., 2005, 2009, 2014). On the other hand, in the genus *Crassula* (Crassulaceae) succulence is also found in species of mesic or wet environments, and has been suggested to be evolutionarily conserved after having been selected in ancestral dry microhabitats (Fradera-Soler *et al*., 2021).

Among epiphytes, monopodial growth evolved from sympodial growth several times independently (Supplementary Data Fig. S2), simultaneously with, or after having evolved, a lateral inflorescence (Supplementary Data Fig. S4), thus allowing the apical bud to grow unlimitedly (Dressler, 1981). Interestingly, all monopodial Oncidiinae and some Angraecinae are twig epiphytes (Chase, 1987; Pridgeon *et al*., 2014), *i*.*e*. they grow on thin branches (< 2.5 cm in diameter), in low or high light zones (Chase, 1987). Twig epiphytes tend to have some specific traits linked to their ephemeral habitat: they are miniature, with short life-cycles (Chase, 1987; Gravendeel *et al*., 2004). All obligate (high light) twig Oncidiinae have a psygmoid (fan-shaped with laterally flattened, unifacial leaves) seedling (Chase, 1987), which is almost identical to the monopodial habit (Chase, 1986). Twigs are indeed ephemeral, oligotrophic habitats with high fluctuations in water availability (Chase, 1986, 1987). Chase (1987) hypothesised that the psygmoid habit could improve the water/surface area relations of these small plants, compared to standard habit. Thus, the monopodial growth of some Oncidiinae could potentially be a functional and morphological convergence to the psygmoid habit, but this is highly speculative as long as the former hypothesis is not tested.

### Succulent traits as key innovations for the diversification of epidendroid orchids

Our comprehensive phylogenetic analysis of the Epidendroideae orchids show that this group appeared during the Eocene climatic optimum, and probably ranged from the Neotropics to Southeast Asia and Australia, possibly via Antarctica (Bush *et al*., 2011). Megathermal forests were then at their most extensive (Bush *et al*., 2011). Stem succulence was found to likely appear in the Eocene 43.1 My ago (95% HPD = 57.3–36.6 Mya) in Southeast Asia to Australia, in the MRCA of Nervilieae and Cymbidieae (Figs. 2 and 3, Supplementary Data Figs. S5 and S6), which is more ancestral than found by Freudenstein and Chase (2015). Confirming the general assumption (Dressler, 1981; Ng and Hew, 2000), the ancestral pseudobulb would have been homoblastic, and the heteroblastic pseudobulb would have then derive from this ancestral homoblastic succulence. The origin of succulence in epidendroid orchids coincides with the origin of Aizoaceae, an entirely African succulent group (41.5 Mya, 95% HPD = 38.7–56.4 Mya) (Arakaki *et al*., 2011; Klak *et al*., 2017). Even though geographical ranges were different between Aizoaceae and Epidendroideae, in the course of the Eocene the global climate gradually became cooler and drier (Bohaty *et al*., 2009; Bush *et al*., 2011), leading to a major retraction of megathermal forests (Bush *et al*., 2011) and to the development of new open habitats (Klak *et al*., 2017). Therefore, since Ringelberg *et al*. (2020) and Anest *et al*. (2021) found that succulence evolved in several groups both in arid desert systems and in semi-arid systems such as savannas, stem succulence in Epidendroideae could have been an adaptation to drier, seasonal forests or savannas.

Stem succulence predated the likely appearances of the epiphytic lifestyle by at least 4.1 My (Fig. 2). Sobralieae lack succulence, but they did not diversify much compared to the other epiphytic groups, with only 304 species (WCSP, 2021) shared out between three genera, of which two contain epiphytic species. Contrary to Freudenstein and Chase (2015), which concluded that ‘there is no key stem modification that appears to be necessary for success as an epiphyte, although thickened leaves are typical’, because ‘[the corm/pseudobulb] is lost in some species-rich groups’, we rather conclude that succulence could have actually been a key innovation (Hunter, 1998) facilitating the transition from terrestrial to epiphytic lifestyle, and preceding the subsequent diversification of epiphytes. Indeed, stem succulence likely appeared earlier than epiphytism and, although it was lost several times in descending epiphytic lineages, it was almost systematically offset by the appearance of leaf succulence, as in Vandeae or Pleurothallidinae (Fig. 2).

To test the impact of succulence traits on the diversification of epiphytes, trait-dependent diversification analyses, for example using Hidden State Speciation and Extinction models (Beaulieu and O’Meara, 2016; Herrera-Alsina *et al*., 2019; Nakov *et al*., 2019), should be carried in the future. However, in order to be accurate these analyses require robust species-level phylogenies, with a good sampling fraction ideally > 40%, which is not the case of the phylogenetic tree we are able to produce until now. Nevertheless, the BAMM diversification analysis carried by Givnish *et al*. (2015) tends to consolidate the hypothesis of succulent traits as key innovations, for they detected a shift towards higher diversification rates in the MRCA of Arethuseae and Cymbidieae, corresponding approximately to the first appearance of epiphytism, and subsequently including the majority of epiphytic Epidendroideae. Stem succulence, by facilitating the transition to the epiphytic lifestyle, would have indirectly enable diversification bursts within the Epidendroideae.

In this study we focused on morphological traits, however physiological traits, notably the water-conserving CAM photosynthesis, could also be important traits contributing to the evolution of epiphytism and the diversification of orchids (Silvera *et al*., 2009; Givnish *et al*., 2015; Silvera and Lasso, 2016). Even though recent studies did not find a link between CAM photosynthesis and an increase in diversification rates, at least in *Bulbophyllum* (Gamisch *et al*., 2021; Hu *et al*., 2022), an estimation of the evolution of CAM photosynthesis has been conducted by Silvera *et al*. (2009) at subtribe and genera levels, then by Givnish *et al*. (2015) at subtribe level, and in spite of high uncertainties in the ancestral state estimations and/or in tree topology, their results nevertheless indicate that CAM photosynthesis could also have appeared prior to epiphytism. In the future, further investigation of the evolutionary history of CAM metabolism in Epidendroideae could thus be interesting.

### Secondary terrestrial genera mostly retained the epiphytic ancestral drought-related traits

Epiphytism has probably been lost at least five times leading to diversification — against three times in Givnish *et al*. (2015), four in Freudenstein and Chase (2015), and six in Chomicki *et al*. (2015)) — during the Oligocene and the Miocene, in different geographical ranges. As reported by Chen *et al*. (2022) in the fern family Polypodiaceae, re-terrestrialization in Calypsoeae likely occurred during the Oligocene in Southeast Asia and the Neotropics, when a glaciation led to the decrease of the area of broad-leaved forests and available habitats, likely increasing the competition in upper canopies (Chen *et al*., 2022). Opening seasonal forests and savannas may have promoted the re-terrestrialization of epiphytes, by allowing more light to reach the floor than broad-leaved forests thus releasing the competition for light in the understorey (Dressler, 1981; Wikström *et al*., 1999), and by allowing plants to occupy a wider range of habitats (Chen *et al*., 2022). In Calypsoeae the terrestrialization probably allowed species to disperse northwards, with almost all genera progressively colonising Eurasia and North America during the Miocene (Fig. 2). Similarly, in Bletiinae the re-terrestrialization (also found by Sosa *et al*. (2016)) probably allowed the simultaneous extension of their geographical range to North America (Figs. 2 and 3).

In Collabieae, Malaxideae, Bletiinae, and Eulophiinae, epiphytism was likely lost during the Miocene. This geological epoch was characterised by fluctuations in the global climate, with rapidly changing environments and temperatures, and by widespread topographic changes which created new habitats (Potter and Szatmari, 2009; Bush *et al*., 2011), enhancing the diversification of many lineages (Potter and Szatmari, 2009). Hence, as hypothesised by Chen *et al*. (2022) for Polypodiaceae, it is likely that new habitats created by climate and topographical changes could have favoured the re-terrestrialization and subsequent diversification of the secondary terrestrial clades in Epidendroideae (Supplementary Data Fig. S10), but also of the epiphytic taxa which seem to have diversified mostly during the Miocene (Supplementary Data Fig. S11). In Eulophiinae, whose diversity is centered on Madagascar (Bone *et al*., 2015), re-terrestrialization occurred between 12 and 13 My ago, i.e. simultaneously with the appearance of the cactiform stem succulence in the genus *Euphorbia* sections *Goniostema, Denisophorbia* and *Deuterocalli* which represent about 70% of the diversity of *Euphorbia* in Madagascar (Aubriot, 2012). The cactiform stem succulence has been found to prevail in areas with seasonal drought but a reliable season of precipitation (Evans *et al*., 2014), which means that seasonal dry forests may have been present in Madagascar at this time and thus could have been favourable to the re-terrestrialization of Eulophiinae.

Interestingly, when looking at the evolutionary history of succulence traits in Epidendroideae, it appears that, on the whole, secondary terrestrial taxa retained their ancestral stem succulence (Fig. 2). In Calypsoeae and Bletiinae, this stem succulence takes the form of underground corms, while in Malaxideae, Collabieae and Eulophiinae it is pseudobulbs (Pridgeon *et al*., 2005, 2009). However, pseudobulbs are sometimes more or less buried, for example in Collabieae genus *Ipsea* (Pridgeon *et al*., 2005; Descourvières, 2011), or in *Eulophia graminea* (Pemberton *et al*., 2008). Actually, burying succulent structures could better protect them from above-ground temperatures extremes and drought, especially in seasonal climates (Pemberton *et al*., 2008; Kumar *et al*., 2022). In addition, Kumar *et al*. (2022) showed that some species in Malaxideae are secondary epiphytes, and that they also retained the perennating organ structure and leaf texture of their parent lineage. This tends to support the hypothesis that seasonal forests could have been the cradle to at least part of both epiphytes and terrestrial Epidendroideae diversity.

The current ecology of secondary terrestrials is also not exactly the same as the primary terrestrial taxa. Indeed, unlike the primary terrestrial taxa, many secondary terrestrials are also found as lithophytes, or even as occasional epiphytes (Pridgeon *et al*., 2005, 2009, 2014). Even though there are differences between the lithophytic and epiphytic habitats, they still share similar rooting conditions (Zotz, 2016), and similar mycorrhizal fungal communities are shared by orchids between the two habitats (Xing *et al*., 2019; Qin *et al*., 2020). Moreover, lithophytic Malaxideae were not found on the rock itself but in the humus-rich and mossy substrate (Hermans *et al*., 2020), resembling some epiphytic conditions. Even among genera that were described as fully terrestrial in Genera Orchidacearum (Pridgeon *et al*., 2005, 2009, 2014), some may not always really root in the soil.

Interestingly, the re-terrestrialization did not particularly lead to a reduction of the number of velamen layers, and this number may have even increased in some secondary terrestrial clades (Supplementary Data Fig. S9). The velamen allows a quick uptake and long retention (increasing with velamen size) of water and nutrients, which is beneficial in environments with low and intermittent water and nutrient supply (Zotz and Winkler, 2013). Other important functions are mechanical protection and, in exposed roots, reduction of heat load (Zotz and Winkler, 2013; Zotz *et al*., 2017) and UV protection (Chomicki *et al*., 2015). Hence, velamentous roots may have been beneficial in secondary terrestrial taxa with lithophytic-like ecology, whose roots can be exposed to a scarce supply of water and UV radiation as for epiphytic taxa. Furthermore, even rooted in soil, numerous taxa in Orchidaceae and other families are velamentous, for the velamen minimises the water loss in dry soils without damaging the root unlike other, nonvelamentous, taxa (Zotz and Winkler, 2013; Zotz *et al*., 2017). Indeed, Zotz *et al*. (2017) found that terrestrial species were most prominent in seasonally dry habitats, which is consistent with our hypothesis of re-terrestrialization events in seasonally dry environments. In addition, Zotz *et al*. (2017) suggested that velamen predated epiphytism, and thus could have been another key innovation (in addition to stem succulence as shown in this study) for the evolution of epiphytism, a view also supported by our results. As well, this is consistent with the hypothesis that epiphytism appeared in mostly seasonally dry climates.

Overall, considering that secondary terrestrials mostly retained their ancestral velamen and succulence traits, we could wonder why some of these taxa are no longer found as epiphytes. The minute orchid seeds should not have constrained dispersal between ground and aerial environments, although different communities of suitable mycorrhizal fungi should (Martos *et al*., 2012). Indeed, given that orchids germinate with mycorrhizal fungi, we hypothesise that these taxa could no longer colonise the epiphytic habitat because of an evolutionary change in their association with mycorrhizal fungal symbionts, which are different between the ground and the trees (Martos *et al*., 2012). Mycorrhizal shifts are also suggested by the fact that all terrestrial lineages derived from epiphytic ancestors have evolved strategies of mycoheterotrophy by parasitizing soil-dwelling, ectomycorrhizal or saprotrophic fungi. Mycoheterotrophy is also often seen as an adaptation to shaded forest understorey (Martos *et al*., 2009). Likewise, even though light conditions of epiphytes span the entire gradient from deep shade to full radiation, many epiphytes may not tolerate shade, and low light in the understorey could thus promote epiphytism (Gravendeel *et al*., 2004; Zotz, 2013). Therefore, because most of Epidendroideae taxa evolved epiphytism, and primary and secondary terrestrial taxa have often evolved mycoheterotrophy, access to light may have been a major constraint in the evolution of Epidendroideae, leading to the evolution of either epiphytism or mycoheterotrophy.

### Assumptions and intrisic limitations of methods

We identified two main methodological limitations intrisic to the methods we used. First, even though our phylogeny is robust and congruent with previous studies based on a higher number of plastid markers, the divergence times were based on only three fossils (including a fossil with an uncertain phylogenetic position), while the Epidendroideae are very much diversified with no less than 23,246 species listed (WCSP, 2021). New orchid fossils would be beneficial to ascertain the divergence times of the subfamily.

Second, models of ancestral state reconstructions rely on several assumptions. For instance discrete state models assumes that transition rates between states are constant throughout the phylogeny, even though hidden states on corHMM alleviate to some extent this assumption. Moreover, all transitions are likely to be slightly underestimated due to the processing of polymorphic traits as uncertainties and not as polymorphisms by corHMM; in other words, we only estimated transitions between sampled genera, but were not able to measure within-genus transitions.

## CONCLUSION

Our analyses show that drought-related traits were likely not adaptations to the epiphytic lifestyle. Rather, epiphytes would have appeared among already drought-adapted terrestrial taxa in seasonally-dry forests, possibly driven by light availability. Particularly, the ancestral stem succulence would have favoured the multiple colonization of aerial environments, and thus would have contributed to the diversification of epiphytes. This character would have evolved once in the form of a homoblastic pseudobulb, later evolving convergently towards heteroblastic pseudobulbs. When lost, stem succulence would have almost always been offset by leaf succulence, indicating that succulent organs could have actually been a key innovation, acting as a precursor enabling the subsequent diversification burst associated with the appearance of the epiphytic lifestyle. However, trait-dependent and trait-independent diversification analyses are still needed to ascertain the diversification patterns of the Epidendroideae and their lifestyles. Other physiological and morpho-anatomical traits, such as the CAM photosynthesis, should also be further investigated in combination with diversification analyses, to further comprehend the macroevolutionary history of epiphytism in Epidendroideae.

