## Supplementary Data for "Gains and losses of the epiphytic lifestyle in epidendroid orchids: review and new analyses with succulence traits"

**RESULTS**

*Estimation of ancestral position of the inflorescence*

Inflorescence was inferred to be ancestrally terminal in Epidendroideae (p = 0.997) (Supplementary Data Fig. S4). Lateral inflorescences appeared convergently several times in the phylogeny in almost all tribes at least at tips (distribution of changes from stochastic mapping: 95% HPD [16, 28], median = 23), and could have been ancestral in (i) the MRCA of Thelasinae 14.9 My ago (p = 0.85, node PP = 1, 95% HPD = 24.2–6.8 Mya), (ii) the MRCA of Collabieae 27.1 My ago (p = 0.54, 95% HPD = 38.2–16.8 Mya), (iii) the MRCA of *Tipularia* and *Changnienia* in Calypsoeae 11.7 My ago (p = 0.85, node PP = 1, 95% HPD = 21.0–4.6 Mya), (iv) the MRCA of *Govenia* and *Aplectrum* in Calypsoeae 21.9 My ago (p = 0.53, node PP = 1, 95% HPD = 30.4–14.7 Mya), (v) the MRCA of Angraecinae and Aeridinae 26.6 My ago (p = 0.87, node PP = 1, 95% HPD = 36.5–19.3 Mya), (vi) the MRCA of Cymbidieae 34.2 My ago (p = 0.81, 95% HPD = 45.1–27.7 Mya). Several reversions to terminal inflorescences occurred independently as well (distribution of changes from stochastic mapping: 95% HPD [2, 8], median = 5).

### TABLES AND FIGURES

#### Table S1

GenBank accession numbers of *matK*, *psaB* and *rbcL* sequences of 203 Epidendroideae genera. The three sequences for each genus originate either from the same voucher (‘voucher’, 69% of genera), from different vouchers but same species (‘sp’, 14%), or from different species (‘x’, 17%). Sequences for the genus *Angraecum* were produced by Givnish *et al.*, 2015 and are provided in their supplementary data, but were not available on GenBank.

| Tribe | Subtribe | Genus | matK | psaB | rbcL | Origin of sequences |
|---|---|---|---|---|---|---|
| ARETHUSEAE | Arethusinae | *Anthogonium* | AF263622.1 | KT885017.1 | AF264153.1 | sp |
| ARETHUSEAE | Arethusinae | *Arethusa* | AF263624.1 | AY380938.1 | AF264154.1 | sp |
| ARETHUSEAE | Arethusinae | *Arundina* | MN171408.1 | MN171408.1 | MN171408.1 | voucher |
| ARETHUSEAE | Arethusinae | *Calopogon* | KJ772600.1 | AY380949.1 | KJ773331.1 | sp |
| ARETHUSEAE | Coelogyninae | *Bletilla* | MT193723.1 | MT193723.1 | MT193723.1 | voucher |
| ARETHUSEAE | Coelogyninae | *Bulleyia* | NC_052739.1 | NC_052739.1 | NC_052739.1 | voucher |
| ARETHUSEAE | Coelogyninae | *Coelogyne* | MT627604.1 | MT627604.1 | MT627604.1 | voucher |
| ARETHUSEAE | Coelogyninae | *Dendrochilum* | MG788099.1 | AY380984.1 | AY381119.1 | sp |
| ARETHUSEAE | Coelogyninae | *Glomera* | AY121742.1 | AY381013.1 | AF074172.1 | x |
| ARETHUSEAE | Coelogyninae | *Panisea* | MT610371.1 | MT610371.1 | MT610371.1 | voucher |
| ARETHUSEAE | Coelogyninae | *Pholidota* | MN879405.1 | MN879405.1 | MN879405.1 | voucher |
| ARETHUSEAE | Coelogyninae | *Pleione* | MK361027.1 | MK361027.1 | MK361027.1 | voucher |
| ARETHUSEAE | Coelogyninae | *Thunia* | MN606292.1 | MN606292.1 | MN606292.1 | voucher |
| CALYPSOEAE | | *Aplectrum* | AF263623.1 | AY380935.1 | AF074108.1 | sp |
| CALYPSOEAE | | *Calypso* | MG874037.1 | MG874037.1 | MG874037.1 | voucher |
| CALYPSOEAE | | *Changnienia* | MN990431.1 | MN990431.1 | MN990431.1 | voucher |
| CALYPSOEAE | | *Corallorhiza* | KM390021.1 | KM390021.1 | KM390021.1 | voucher |
| CALYPSOEAE | | *Cremastra* | MN990434.1 | MN990434.1 | MN990434.1 | voucher |
| CALYPSOEAE | | *Dactylostalix* | KM526761.1 | KM526760.1 | KM526772.1 | voucher |
| CALYPSOEAE | | *Govenia* | AF263664.1 | AY381017.1 | AF074175.1 | x |
| CALYPSOEAE | | *Oreorchis* | MN990443.1 | MN990443.1 | MN990443.1 | voucher |
| CALYPSOEAE | | *Tipularia* | MN990442.1 | MN990442.1 | MN990442.1 | voucher |
| COLLABIEAE | | *Acanthophippium* | AF263618.1 | AY380927.1 | AF074100.1 | voucher |

| | | | | | | |
|---|---|---|---|---|---|---|
| COLLABIEAE | | *Calanthe* | KF753635.1 | KF753635.1 | KF753635.1 | voucher |
| COLLABIEAE | | *Cephalantheropsis* | NC_050868.1 | NC_050868.1 | NC_050868.1 | voucher |
| COLLABIEAE | | *Chrysoglossum* | KF852687.1 | KF852620.1 | KF852732.1 | voucher |
| COLLABIEAE | | *Collabium* | KF852672.1 | KF852604.1 | KF852716.1 | voucher |
| COLLABIEAE | | *Eriodes* | KF852674.1 | KF852606.1 | KF852718.1 | voucher |
| COLLABIEAE | | *Hancockia* | KF852675.1 | KF852607.1 | KF852719.1 | voucher |
| COLLABIEAE | | *Nephelaphyllum* | KF852689.1 | KF852622.1 | KF852734.1 | voucher |
| COLLABIEAE | | *Phaius* | MN708349.1 | MN708349.1 | MN708349.1 | voucher |
| COLLABIEAE | | *Risleya* | KF852685.1 | KF852617.1 | KF852729.1 | voucher |
| COLLABIEAE | | *Spathoglottis* | AY368429.1 | AY381077.1 | AY381134.1 | x |
| COLLABIEAE | | *Tainia* | NC_045862.1 | NC_045862.1 | NC_045862.1 | voucher |
| CYMBIDIEAE | Catasetinae | *Catasetum* | AF263637.1 | AY380951.1 | AF074121.1 | x |
| CYMBIDIEAE | Catasetinae | *Cycnoches* | AY368401.1 | KT750206.1 | AY368355.1 | x |
| CYMBIDIEAE | Catasetinae | *Dressleria* | AY368406.1 | AY380991.1 | AF074153.1 | x |
| CYMBIDIEAE | Catasetinae | *Galeandra* | AY368408.1 | AY381011.1 | AF074171.1 | voucher |
| CYMBIDIEAE | Catasetinae | *Mormodes* | AY368417.1 | KT750211.1 | KT750221.1 | x |
| CYMBIDIEAE | Coeliopsidinae | *Lycomormium* | AY368414.1 | AY381029.1 | AF074186.1 | x |
| CYMBIDIEAE | Coeliopsidinae | *Peristeria* | MF349960.1 | KT750213.1 | KT750222.1 | sp |
| CYMBIDIEAE | Cymbidiinae | *Acriopsis* | EF065575.1 | KT750204.1 | KT750215.1 | x |
| CYMBIDIEAE | Cymbidiinae | *Ansellia* | KT750199.1 | KT750205.1 | KT750216.1 | voucher |
| CYMBIDIEAE | Cymbidiinae | *Cymbidium* | MT576628.1 | MT576628.1 | MT576628.1 | voucher |
| CYMBIDIEAE | Cymbidiinae | *Dipodium* | MN200386.1 | MN200386.1 | MN200386.1 | voucher |
| CYMBIDIEAE | Cymbidiinae | *Grammatophyllum* | AF470458.1 | AY381018.1 | AF074176.1 | x |
| CYMBIDIEAE | Cymbidiinae | *Thecopus* | KT750203.1 | KT750214.1 | KT750223.1 | voucher |
| CYMBIDIEAE | Cyrtopodiinae | *Cyrtopodium* | KT750200.1 | KT750207.1 | KT750217.1 | voucher |
| CYMBIDIEAE | Eriopsidinae | *Eriopsis* | DQ210866.1 | AY381007.1 | AF074167.1 | sp |
| CYMBIDIEAE | Eulophiinae | *Acrolophia* | KF358114.1 | | KF358051.1 | voucher |
| CYMBIDIEAE | Eulophiinae | *Eulophia* | MG181954.1 | MG181954.1 | MG181954.1 | voucher |
| CYMBIDIEAE | Eulophiinae | *Eulophiella* | FR832765.1 | | FN870819.1 | voucher |
| CYMBIDIEAE | Eulophiinae | *Geodorum* | MT153204.1 | MT153204.1 | MT153204.1 | voucher |
| CYMBIDIEAE | Eulophiinae | *Grammangis* | KT750202.1 | KT750210.1 | KT750219.1 | voucher |
| CYMBIDIEAE | Eulophiinae | *Oeceoclades* | KF358118.1 | KT750212.1 | KF358008.1 | sp |
| CYMBIDIEAE | Eulophiinae | *Paralophia* | LN831620.1 | | LN831611.1 | voucher |
| CYMBIDIEAE | Maxillariinae | *Bifrenaria* | EF065567.1 | AY380941.1 | AF074112.1 | sp |
| CYMBIDIEAE | Maxillariinae | *Lycaste* | MH748890.1 | AY381028.1 | MH748851.1 | sp |
| CYMBIDIEAE | Maxillariinae | *Maxillaria* | DQ209871.1 | AY381033.1 | MN719148.1 | x |
| CYMBIDIEAE | Maxillariinae | *Neomoorea* | AF239437.1 | AY381042.1 | AF074198.1 | sp |
| CYMBIDIEAE | Maxillariinae | *Rudolfiella* | FJ564977.1 | FJ534334.1 | FJ534212.1 | voucher |
| CYMBIDIEAE | Maxillariinae | *Xylobium* | MH748945.1 | AY381097.1 | MH749097.1 | x |

| | | | | | | |
|---|---|---|---|---|---|---|
| CYMBIDIEAE | Oncidiinae | *Aspasia* | FJ563860.1 | FJ534283.1 | FJ534160.1 | voucher |
| CYMBIDIEAE | Oncidiinae | *Brassia* | AF350601.1 | FJ534314.1 | FJ534192.1 | voucher |
| CYMBIDIEAE | Oncidiinae | *Capanemia* | FJ563840.1 | FJ534262.1 | FJ534139.1 | voucher |
| CYMBIDIEAE | Oncidiinae | *Caucaea* | FJ565012.1 | FJ534343.1 | FJ534221.1 | voucher |
| CYMBIDIEAE | Oncidiinae | *Chytroglossa* | FJ565112.1 | FJ534366.1 | FJ534244.1 | voucher |
| CYMBIDIEAE | Oncidiinae | *Cischweinfia* | FJ565125.1 | FJ534370.1 | FJ534248.1 | voucher |
| CYMBIDIEAE | Oncidiinae | *Comparettia* | FJ565090.1 | FJ534359.1 | FJ534237.1 | voucher |
| CYMBIDIEAE | Oncidiinae | *Cuitlauzina* | FJ565149.1 | FJ534374.1 | FJ534253.1 | voucher |
| CYMBIDIEAE | Oncidiinae | *Cyrtochiloides* | AF433009.1 | FJ534273.1 | FJ534150.1 | voucher |
| CYMBIDIEAE | Oncidiinae | *Cyrtochilum* | FJ564969.1 | FJ534331.1 | FJ534209.1 | voucher |
| CYMBIDIEAE | Oncidiinae | *Eloyella* | DQ315888.1 | FJ534327.1 | FJ534205.1 | voucher |
| CYMBIDIEAE | Oncidiinae | *Erycina* | JF746994.1 | JF746994.1 | JF746994.1 | voucher |
| CYMBIDIEAE | Oncidiinae | *Fernandezia* | FJ565010.1 | FJ534341.1 | FJ534219.1 | voucher |
| CYMBIDIEAE | Oncidiinae | *Gomesa* | AF350632.1 | FJ534277.1 | FJ534154.1 | voucher |
| CYMBIDIEAE | Oncidiinae | *Grandiphyllum* | FJ564946.1 | FJ534321.1 | FJ534199.1 | voucher |
| CYMBIDIEAE | Oncidiinae | *Hintonella* | FJ564940.1 | FJ534317.1 | FJ534195.1 | voucher |
| CYMBIDIEAE | Oncidiinae | *Hofmeisterella* | FJ565091.1 | FJ534360.1 | FJ534238.1 | voucher |
| CYMBIDIEAE | Oncidiinae | *Ionopsis* | FJ565042.1 | FJ534347.1 | FJ534225.1 | voucher |
| CYMBIDIEAE | Oncidiinae | *Leochilus* | FJ563848.1 | FJ534272.1 | FJ534149.1 | voucher |
| CYMBIDIEAE | Oncidiinae | *Lockhartia* | FJ565011.1 | FJ534342.1 | FJ534220.1 | voucher |
| CYMBIDIEAE | Oncidiinae | *Macradenia* | FJ564839.1 | FJ534303.1 | FJ534181.1 | voucher |
| CYMBIDIEAE | Oncidiinae | *Macroclinium* | FJ564931.1 | FJ534313.1 | FJ534191.1 | voucher |
| CYMBIDIEAE | Oncidiinae | *Miltonia* | FJ565115.1 | FJ534367.1 | FJ534245.1 | voucher |
| CYMBIDIEAE | Oncidiinae | *Nohawilliamsia* | FJ563950.1 | FJ534311.1 | FJ534189.1 | voucher |
| CYMBIDIEAE | Oncidiinae | *Notylia* | FJ564961.1 | FJ534330.1 | FJ534208.1 | voucher |
| CYMBIDIEAE | Oncidiinae | *Notyliopsis* | FJ565086.1 | FJ534356.1 | FJ534234.1 | voucher |
| CYMBIDIEAE | Oncidiinae | *Oliveriana* | FJ564814.1 | FJ534296.1 | FJ534174.1 | voucher |
| CYMBIDIEAE | Oncidiinae | *Oncidium* | KM032624.1 | KM032624.1 | KM032624.1 | voucher |
| CYMBIDIEAE | Oncidiinae | *Ornithocephalus* | FJ565049.1 | FJ534350.1 | FJ534228.1 | voucher |
| CYMBIDIEAE | Oncidiinae | *Otoglossum* | FJ565100.1 | FJ534363.1 | FJ534241.1 | voucher |
| CYMBIDIEAE | Oncidiinae | *Phymatidium* | FJ563942.1 | FJ534305.1 | FJ534183.1 | voucher |
| CYMBIDIEAE | Oncidiinae | *Plectrophora* | FJ564979.1 | FJ534335.1 | FJ534213.1 | voucher |
| CYMBIDIEAE | Oncidiinae | *Polyotidium* | FJ563960.1 | FJ534323.1 | FJ534201.1 | voucher |
| CYMBIDIEAE | Oncidiinae | *Psychopsiella* | FJ565152.1 | FJ534375.1 | FJ534254.1 | voucher |
| CYMBIDIEAE | Oncidiinae | *Psychopsis* | FJ564712.1 | FJ534264.1 | FJ534141.1 | voucher |
| CYMBIDIEAE | Oncidiinae | *Pterostemma* | FJ563948.1 | FJ534310.1 | FJ534188.1 | voucher |
| CYMBIDIEAE | Oncidiinae | *Rhynchostele* | AF350609.1 | FJ534280.1 | FJ534157.1 | voucher |
| CYMBIDIEAE | Oncidiinae | *Rodriguezia* | FJ564975.1 | FJ534333.1 | FJ534211.1 | voucher |
| CYMBIDIEAE | Oncidiinae | *Rossioglossum* | FJ563837.1 | FJ534258.1 | FJ534136.1 | voucher |

| | | | | | | |
|---|---|---|---|---|---|---|
| CYMBIDIEAE | Oncidiinae | *Saundersia* | FJ564734.1 | FJ534278.1 | FJ534155.1 | voucher |
| CYMBIDIEAE | Oncidiinae | *Schunkea* | FJ563933.1 | FJ534300.1 | FJ534178.1 | voucher |
| CYMBIDIEAE | Oncidiinae | *Seegeriella* | FJ564829.1 | FJ534299.1 | FJ534177.1 | voucher |
| CYMBIDIEAE | Oncidiinae | *Solenidium* | FJ564956.1 | FJ534328.1 | FJ534206.1 | voucher |
| CYMBIDIEAE | Oncidiinae | *Sutrina* | FJ564828.1 | FJ534298.1 | FJ534176.1 | voucher |
| CYMBIDIEAE | Oncidiinae | *Systeloglossum* | AF350607.1 | FJ534287.1 | FJ534165.1 | sp |
| CYMBIDIEAE | Oncidiinae | *Telipogon* | AF239488.1 | AY381080.1 | AF074231.1 | voucher |
| CYMBIDIEAE | Oncidiinae | *Tolumnia* | AF350611.1 | FJ534281.1 | FJ534158.1 | voucher |
| CYMBIDIEAE | Oncidiinae | *Trichocentrum* | FJ565028.1 | FJ534345.1 | FJ534223.1 | voucher |
| CYMBIDIEAE | Oncidiinae | *Trichoceros* | FJ564953.1 | FJ534348.1 | FJ534226.1 | sp |
| CYMBIDIEAE | Oncidiinae | *Trichopilia* | FJ565053.1 | FJ534351.1 | FJ534229.1 | voucher |
| CYMBIDIEAE | Oncidiinae | *Trizeuxis* | FJ563850.1 | FJ534274.1 | FJ534151.1 | voucher |
| CYMBIDIEAE | Oncidiinae | *Vitekorchis* | FJ565094.1 | FJ534362.1 | FJ534240.1 | voucher |
| CYMBIDIEAE | Oncidiinae | *Warmingia* | FJ563944.1 | FJ534306.1 | FJ534184.1 | voucher |
| CYMBIDIEAE | Oncidiinae | *Zelenkoa* | FJ564942.1 | FJ534318.1 | FJ534196.1 | voucher |
| CYMBIDIEAE | Oncidiinae | *Zygostates* | FJ565111.1 | FJ534365.1 | FJ534243.1 | voucher |
| CYMBIDIEAE | Stanhopeinae | *Acineta* | AF263619.1 | AY380929.1 | AF074102.1 | sp |
| CYMBIDIEAE | Stanhopeinae | *Coryanthes* | AY368398.1 | AY380969.1 | AF074134.1 | x |
| CYMBIDIEAE | Stanhopeinae | *Houlletia* | AF239467.1 | AY381020.1 | AF074178.1 | sp |
| CYMBIDIEAE | Stanhopeinae | *Kegeliella* | AF263666.1 | AY381024.1 | AF074181.1 | x |
| CYMBIDIEAE | Stanhopeinae | *Stanhopea* | AF239445.1 | AY381079.1 | MN719156.1 | x |
| CYMBIDIEAE | Zygopetalinae | *Chaubardia* | AF239419.1 | AY381021.1 | AF074179.1 | sp |
| CYMBIDIEAE | Zygopetalinae | *Cryptarrhena* | AY368399.1 | AY380974.1 | AF074138.1 | x |
| CYMBIDIEAE | Zygopetalinae | *Dichaea* | EU123657.1 | AY380986.1 | AF074149.1 | x |
| CYMBIDIEAE | Zygopetalinae | *Zygopetalum* | AF263689.1 | AY381098.1 | AF074246.1 | x |
| DENDROBIEAE | | *Bulbophyllum* | MN737573.1 | MN737573.1 | MN737573.1 | voucher |
| DENDROBIEAE | | *Cadetia* | EF079346.1 | AY380945.1 | D58406.1 | x |
| DENDROBIEAE | | *Dendrobium* | LC192954.1 | LC192954.1 | LC192954.1 | voucher |
| DENDROBIEAE | | *Epigeneium* | KF143738.1 | AY380997.1 | KF177669.1 | x |
| EPIDENDREAE | Bletiinae | *Basiphyllaea* | MK726085.1 | MK726085.1 | MK726085.1 | voucher |
| EPIDENDREAE | Bletiinae | *Bletia* | MK726066.1 | MK726066.1 | MK726066.1 | voucher |
| EPIDENDREAE | Bletiinae | *Hexalectris* | MK726072.1 | MK726072.1 | MK726072.1 | voucher |
| EPIDENDREAE | Chysinae | *Chysis* | AF263640.1 | AY380956.1 | AF074126.1 | sp |
| EPIDENDREAE | Coeliinae | *Coelia* | AF263631.1 | AY380966.1 | AF264158.1 | sp |
| EPIDENDREAE | Laeliinae | *Arpophyllum* | AF263625.1 | AY380939.1 | AF074110.1 | sp |
| EPIDENDREAE | Laeliinae | *Cattleya* | KP168671.1 | KP168671.1 | KP168671.1 | voucher |
| EPIDENDREAE | Laeliinae | *Epidendrum* | KJ472367.1 | AY380996.1 | MN719140.1 | sp |
| EPIDENDREAE | Laeliinae | *Meiracyllium* | AF263670.1 | AY381037.1 | AF074192.1 | sp |
| EPIDENDREAE | Laeliinae | *Prosthechea* | AY396126.1 | AY380995.1 | KJ773788.1 | sp |

| | | | | | | |
|---|---|---|---|---|---|---|
| EPIDENDREAE | Pleurothallidinae | *Anathallis* | MH979332.1 | MH979332.1 | MH979332.1 | voucher |
| EPIDENDREAE | Pleurothallidinae | *Dilomilis* | AY368404.1 | AY380987.1 | AF074150.1 | voucher |
| EPIDENDREAE | Pleurothallidinae | *Masdevallia* | KP205432.1 | KP205432.1 | KP205432.1 | voucher |
| EPIDENDREAE | Pleurothallidinae | *Pleurothallis* | AF263680.1 | AY381059.1 | AF264174.1 | x |
| EPIDENDREAE | Pleurothallidinae | *Restrepia* | AY370654.1 | AY381072.1 | AY370653.1 | x |
| MALAXIDEAE | | *Dienia* | KF852679.1 | KF852611.1 | KF852723.1 | voucher |
| MALAXIDEAE | | *Liparis* | MN627759.1 | MN627759.1 | MN627759.1 | voucher |
| MALAXIDEAE | | *Malaxis* | KF852680.1 | KF852612.1 | KF852724.1 | voucher |
| MALAXIDEAE | | *Oberonia* | KX871235.1 | KX871235.1 | KX871235.1 | voucher |
| MALAXIDEAE | | *Oberonioides* | MT559316.1 | MT559316.1 | MT559316.1 | voucher |
| NEOTTIEAE | | *Aphyllorchis* | NC_030703.1 | NC_030703.1 | NC_030703.1 | voucher |
| NEOTTIEAE | | *Cephalanthera* | MH590346.1 | MH590346.1 | MH590346.1 | voucher |
| NEOTTIEAE | | *Epipactis* | KU551267.1 | KU551267.1 | KU551267.1 | voucher |
| NEOTTIEAE | | *Neottia* | KU551271.1 | KU551271.1 | KU551271.1 | voucher |
| NEOTTIEAE | | *Palmorchis* | MH590357.1 | MH590357.1 | MH590357.1 | voucher |
| NERVILIEAE | Nerviliinae | *Nervilia* | JN004498.1 | EF089166.1 | JN005559.1 | sp |
| PODOCHILEAE | Eriinae | *Appendicula* | KY239521.1 | KT885018.1 | AF518050.1 | sp |
| PODOCHILEAE | Eriinae | *Ceratostylis* | KY239523.1 | KT885020.1 | AY368353.1 | x |
| PODOCHILEAE | Eriinae | *Eria* | MN477202.1 | MN477202.1 | MN477202.1 | voucher |
| PODOCHILEAE | Eriinae | *Oxystophyllum* | KF143742.1 | KT885028.1 | KF177673.1 | sp |
| PODOCHILEAE | Eriinae | *Podochilus* | KY239515.1 | AY381060.1 | AF074218.1 | voucher |
| PODOCHILEAE | Eriinae | *Trichotosia* | AY368432.1 | AY381085.1 | AF074235.1 | voucher |
| PODOCHILEAE | Thelasinae | *Phreatia* | KY239494.1 | AY381056.1 | AF074214.1 | x |
| PODOCHILEAE | Thelasinae | *Thelasis* | KY239490.1 | KT885035.1 | AF518047.1 | x |
| SOBRALIEAE | | *Elleanthus* | KR260986.1 | KR260986.1 | KR260986.1 | voucher |
| SOBRALIEAE | | *Sobralia* | KM032623.1 | KM032623.1 | KM032623.1 | voucher |
| TRIPHOREAE | Triphorinae | *Monophyllorchis* | EF065603.1 | AY381040.1 | AF074195.1 | x |
| TRIPHOREAE | Triphorinae | *Triphora* | AY368433.1 | AY381086.1 | AF074236.1 | voucher |
| TROPIDIEAE | | *Corymborkis* | AF263647.1 | AY380972.1 | AF074136.1 | x |
| TROPIDIEAE | | *Tropidia* | AJ310078.1 | AY381087.1 | AF074237.1 | x |
| VANDEAE | Aeridinae | *Acampe* | MN124419.1 | MN124419.1 | MN124419.1 | voucher |
| VANDEAE | Aeridinae | *Aerides* | EF655773.1 | KT885015.1 | MT919285.1 | sp |
| VANDEAE | Aeridinae | *Arachnis* | KJ733545.1 | KT885019.1 | JQ933223.1 | x |
| VANDEAE | Aeridinae | *Cleisomeria* | MN124442.1 | MN124442.1 | MN124442.1 | voucher |
| VANDEAE | Aeridinae | *Cleisostoma* | MN124421.1 | MN124421.1 | MN124421.1 | voucher |
| VANDEAE | Aeridinae | *Diploprora* | MN124409.1 | MN124409.1 | MN124409.1 | voucher |
| VANDEAE | Aeridinae | *Gastrochilus* | NC_035833.1 | NC_035833.1 | NC_035833.1 | voucher |
| VANDEAE | Aeridinae | *Grosourdya* | KJ733578.1 | KT885022.1 | KT884994.1 | voucher |
| VANDEAE | Aeridinae | *Holcoglossum* | MK442935.1 | MK442935.1 | MK442935.1 | voucher |

| | | | | | | |
|---|---|---|---|---|---|---|
| VANDEAE | Aeridinae | *Luisia* | KJ733581.1 | KT885024.1 | KT884995.1 | voucher |
| VANDEAE | Aeridinae | *Papilionanthe* | KC823036.1 | KT885029.1 | FN870889.1 | x |
| VANDEAE | Aeridinae | *Pelatantheria* | KX871232.1 | KX871232.1 | KX871232.1 | voucher |
| VANDEAE | Aeridinae | *Phalaenopsis* | MT822270.1 | MT822270.1 | MT822270.1 | voucher |
| VANDEAE | Aeridinae | *Pomatocalpa* | MN124411.1 | MN124411.1 | MN124411.1 | voucher |
| VANDEAE | Aeridinae | *Renanthera* | KJ733597.1 | KT885030.1 | FN870918.1 | x |
| VANDEAE | Aeridinae | *Rhynchostylis* | EF655804.1 | KT885031.1 | KX527029.1 | sp |
| VANDEAE | Aeridinae | *Robiquetia* | MN124410.1 | MN124410.1 | MN124410.1 | voucher |
| VANDEAE | Aeridinae | *Sarcoglyphis* | MN124408.1 | MN124408.1 | MN124408.1 | voucher |
| VANDEAE | Aeridinae | *Schoenorchis* | MN124407.1 | MN124407.1 | MN124407.1 | voucher |
| VANDEAE | Aeridinae | *Smitinandia* | MN124406.1 | MN124406.1 | MN124406.1 | voucher |
| VANDEAE | Aeridinae | *Stereochilus* | MN124431.1 | MN124431.1 | MN124431.1 | voucher |
| VANDEAE | Aeridinae | *Thrixspermum* | MN725094.1 | MN725094.1 | MN725094.1 | voucher |
| VANDEAE | Aeridinae | *Trichoglottis* | MN124432.1 | MN124432.1 | MN124432.1 | voucher |
| VANDEAE | Aeridinae | *Uncifera* | MN124433.1 | MN124433.1 | MN124433.1 | voucher |
| VANDEAE | Aeridinae | *Vanda* | NC_048458.1 | NC_048458.1 | NC_048458.1 | voucher |
| VANDEAE | Aeridinae | *Vandopsis* | MN124443.1 | MN124443.1 | MN124443.1 | voucher |
| VANDEAE | Agrostophyllinae | *Agrostophyllum* | MN990432.1 | MN990432.1 | MN990432.1 | voucher |
| VANDEAE | Agrostophyllinae | *Earina* | AF263656.1 | AY380993.1 | AF074155.1 | x |
| VANDEAE | Angraecinae | *Aerangis* | MK685600.1 | AY380931.1 | KU748383.1 | sp |
| VANDEAE | Angraecinae | *Aeranthes* | KJ021021.1 | AY380932.1 | AF074104.1 | x |
| VANDEAE | Angraecinae | *Angraecum* | Givnish et al., 2015 | Givnish et al., 2015 | Givnish et al., 2015 | voucher |
| VANDEAE | Angraecinae | *Rhipidoglossum* | MK685650.1 | AY380985.1 | AF074147.1 | sp |
| VANDEAE | Polystachyinae | *Polystachya* | AY368426.1 | AY381064.1 | AF074222.1 | x |

**Table S2**

Time-stratified dispersal multiplers between bioregions applied to BioGeoBEARS models, following Givnish *et al*., 2016. N: North America, T: Neotropics, E: Eurasia, F: Africa, S: Southeast Asia, A: Australia, P: Pacific.

0-2 Mya

| | N | T | E | F | S | A | P |
|---|---|---|---|---|---|---|---|
| N | 1 | 1 | 0.1 | 0.1 | 0.1 | 0.1 | 0.1 |
| T | 1 | 1 | 0.1 | 0.1 | 0.1 | 0.1 | 0.1 |
| E | 0.1 | 0.1 | 1 | 1 | 0.5 | 0.1 | 0.1 |
| F | 0.1 | 0.1 | 1 | 1 | 0.1 | 0.1 | 0.1 |
| S | 0.1 | 0.1 | 0.5 | 0.1 | 1 | 0.5 | 0.1 |
| A | 0.1 | 0.1 | 0.1 | 0.1 | 0.5 | 1 | 0.5 |
| P | 0.1 | 0.1 | 0.1 | 0.1 | 0.1 | 0.5 | 1 |

2-30 Mya

| | N | T | E | F | S | A | P |
|---|---|---|---|---|---|---|---|
| N | 1 | 0.5 | 0.1 | 0.1 | 0.1 | 0.1 | 0.1 |
| T | 0.5 | 1 | 0.1 | 0.1 | 0.1 | 0.1 | 0.1 |
| E | 0.1 | 0.1 | 1 | 1 | 0.5 | 0.1 | 0.1 |
| F | 0.1 | 0.1 | 1 | 1 | 0.1 | 0.1 | 0.1 |
| S | 0.1 | 0.1 | 0.5 | 0.1 | 1 | 0.1 | 0.1 |
| A | 0.1 | 0.1 | 0.1 | 0.1 | 0.1 | 1 | 0.5 |
| P | 0.1 | 0.1 | 0.1 | 0.1 | 0.1 | 0.5 | 1 |

30-60 Mya

| | N | T | E | F | S | A | P |
|---|---|---|---|---|---|---|---|
| N | 1 | 0.5 | 0.5 | 0.1 | 0.1 | 0.1 | 0.1 |
| T | 0.5 | 1 | 0.1 | 0.1 | 0.1 | 0.1 | 0.1 |
| E | 0.5 | 0.1 | 1 | 1 | 0.5 | 0.1 | 0.1 |
| F | 0.1 | 0.1 | 1 | 1 | 0.1 | 0.1 | 0.1 |
| S | 0.1 | 0.1 | 0.5 | 0.1 | 1 | 0.1 | 0.1 |
| A | 0.1 | 0.1 | 0.1 | 0.1 | 0.1 | 1 | 0.5 |
| P | 0.1 | 0.1 | 0.1 | 0.1 | 0.1 | 0.5 | 1 |

**Table S3**

Symmetric (SYM) transition rate matrices between combined states used in corHMM for testing the correlation between lifestyle and binary morphological traits (stem succulence, leaf succulence, growth form, position of inflorescence). Here the test for the correlated evolution of lifestyle and growth form is shown as an example. Numbers indicate the transition rate category, with 0 meaning that transitions are not allowed between corresponding combined states. The decorrelated matrix allows to move from one state to the other inside a trait, whatever the state of the other trait is. For example, rate category 1 allows transitions to occur at the same rate between the epiphytic and the terrestrial lifestyles, independently of the growth form being monopodial or sympodial.

Correlated SYM matrix

| | Epiphyte & Monopodial | Epiphyte & Sympodial | Terrestrial & Monopodial | Terrestrial & Sympodial |
|---|---|---|---|---|
| Epiphyte & Monopodial | 0 | 1 | 2 | 0 |
| Epiphyte & Sympodial | 1 | 0 | 0 | 3 |
| Terrestrial & Monopodial | 2 | 0 | 0 | 4 |
| Terrestrial & Sympodial | 0 | 3 | 4 | 0 |

Decorrelated SYM matrix

| | Epiphyte & Monopodial | Epiphyte & Sympodial | Terrestrial & Monopodial | Terrestrial & Sympodial |
|---|---|---|---|---|
| Epiphyte & Monopodial | 0 | 3 | 4 | 0 |
| Epiphyte & Sympodial | 2 | 0 | 0 | 4 |
| Terrestrial & Monopodial | 1 | 0 | 0 | 3 |
| Terrestrial & Sympodial | 0 | 1 | 2 | 0 |

**Table S4**

Survey of morphological characteristics and lifestyle of the 201 accepted genera of Epidendroideae sampled in the phylogeny. Two synonymised genera (*Epigeneium* and *Cadetia*, synonym: *Dendrobium*) included in the phylogeny were excluded from this table. Data compiled from *Genera Orchidacearum* (Pridgeon *et al.*, 2005, 2009, 2014). Lithophytes were merged with epiphytes or terrestrial genera, no genus having only a litophytic lifestyle. Stem succulence includes corms and pseudobulbs. Leaf succulence includes leaves described as coriaceous or fleshy.

| Trait | State | Accepted genera (201) | % |
|---|---|---|---|
| Lifestyle | Epiphyte | 126 | 62.7 |
| | Terrestrial | 50 | 24.9 |
| | Epiphyte & Terrestrial | 25 | 12.4 |
| Stem succulence | Absence | 75 | 37.3 |
| | Succulent | 104 | 51.7 |
| | Absence & Succulent | 22 | 10.9 |
| Leaf succulence | Absence | 126 | 62.7 |
| | Succulent | 74 | 36.8 |
| | Absence & Succulent | 21 | 10.4 |
| Position of inflorescence | Lateral | 143 | 71.1 |
| | Terminal | 48 | 23.9 |
| | Lateral & Terminal | 10 | 5 |
| Growth form | Monopodial | 37 | 18.4 |
| | Sympodial | 160 | 79.6 |
| | Monopodial & Sympodial | 4 | 2 |

**Table S5**

AICc comparison of BioGeoBEARS models. d, e, j: values for dispersal, extinction, and founder-event speciation parameters, respectively. delta AICc: difference of AICc between the model and the model with minimal AICc. AICc weight: proportion of the total predictive power of all models explained by the model.

| BioGeoBEARS model | Number of parameters | d | e | j | AIC | AICc | delta AICc | AICc weight |
|---|---|---|---|---|---|---|---|---|
| BayAreaLike*+J | 3 | 0.02 | 0.02 | 0.02 | 887.92 | 888.04 | 0 | 0.72 |
| BayAreaLike* | 2 | 0.02 | 0.02 | 0 | 889.89 | 889.95 | 1.91 | 0.28 |
| DEC* | 2 | 0.04 | 0 | 0 | 973.51 | 973.57 | 85.53 | 0 |
| DEC*+J | 3 | 0.04 | 0 | 0 | 975.51 | 975.63 | 87.59 | 0 |
| DIVALIKE* | 2 | 0.05 | 0.01 | 0 | 1004.1 | 1004.16 | 116.12 | 0 |
| DIVALIKE*+J | 3 | 0.05 | 0.01 | 0 | 1006.1 | 1006.22 | 118.18 | 0 |

**Table S6**

Likelihood ratio-tests for nested BioGeoBEARS models. LnL alt, LnL null: log-likelihood of the data given the alternative or null model, respectively. DF alt, DF null: degrees of freedom of alternative and null models, respectively. p-value: p-value of the one-tailed chi-squared test.

| Alternative model (alt) | Null model (null) | LnL alt | LnL null | DF alt | DF null | D statistic | p-value |
|---|---|---|---|---|---|---|---|
| DEC*+J | DEC* | -484.8 | -484.8 | 3 | 2 | -0.0039 | 1 |
| DIVALIKE*+J | DIVALIKE* | -500.1 | -500 | 3 | 2 | -0.0032 | 1 |
| BAYAREALIKE*+J | BAYAREALIKE* | -441 | -442.9 | 3 | 2 | 3.97 | 0.046 |

**Figure S1**

Evolution of lifestyle in epidendroid orchids. (A) Dated phylogenetic tree reconstructed from 203 genera in 14 tribes of epidendroid orchids. Branches with posterior probability < 0.95 are coloured coral-red. Nodes that were constrained in the analysis are indicated by stars (in red when the quartet support for the main topology was < 50% in the coalescent-species tree of Pérez-Escobar *et al.* 2021). (B) and (C) Estimation of ancestral states in the Epidendroideae genus tree with corHMM. Pie charts at nodes represent ancestral lifestyles and their probabilities. The character states of the present genera are represented on the right side of the tree by coloured boxes. In addition, genera with mycoheterotrophic species are indicated by a mushroom symbol. Cymbidiinae are split into three clades which are indicated by one, two and three stars after the subtribe name.

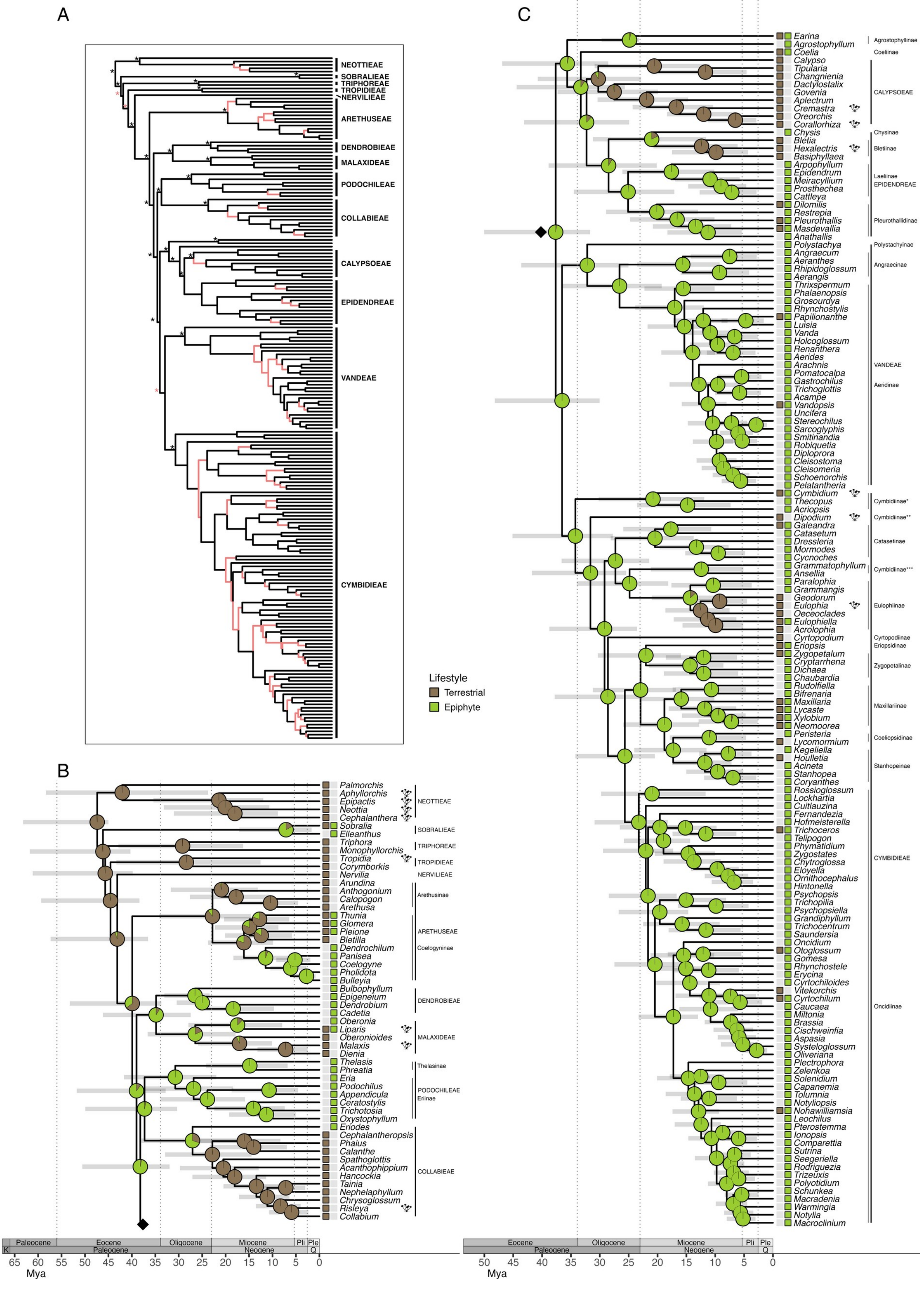
A
NEOTTIEAE
SOBRALIEAE
TRIPHOREAE
TROPIDIEAE
NERVILIEAE
ARETHUSEAE
DENDROBIEAE
MALAXIDEAE
PODOCHILEAE
COLLABIEAE
CALYPSOEAE
EPIDENDREAE
VANDEAE
CYMBIDIEAE
Lifestyle
Terrestrial
Epiphyte
B
Palmorchis
Aphyllorchis
Epipactis
Neottia
Cephalanthera
Sobralia
Elleanthus
Triphora
Monophyllorchis
Tropidia
Corymborkis
Nervilia
Arundina
Anthogonium
Calopogon
Arethusa
Thunia
Glomera
Pleione
Bletilla
Dendrochilum
Panisea
Coelogyne
Pholidota
Bulleyia
Bulbophyllum
Epigeneium
Dendrobium
Cadetia
Oberonia
Liparis
Oberonioides
Malaxis
Dienia
Thelasis
Phreatia
Eria
Podochilus
Appendicula
Ceratostylis
Trichotosia
Oxystophyllum
Eriodes
Cephalanthropsis
Phaius
Calanthe
Spathoglottis
Acanthophippium
Hancockia
Tainia
Nephelaphyllum
Chrysoglossum
Risleya
Collabium
NEOTTIEAE
SOBRALIEAE
TRIPHOREAE
TROPIDIEAE
NERVILIEAE
Arethusinae
ARETHUSEAE
Coelogyninae
DENDROBIEAE
MALAXIDEAE
Thelasinae
PODOCHILEAE
Eriinae
COLLABIEAE
Paleocene
Eocene
Oligocene
Miocene
Pli
Ple
K
Paleogene
Neogene
Q
65 60 55 50 45 40 35 30 25 20 15 10 5 0
Mya
C
Earina
Agrostophyllum
Coelia
Calypso
Tipularia
Changnienia
Dactylostalix
Govenia
Aplectrum
Cremastra
Oreorchis
Corallorhiza
Chysis
Bletia
Hexalectris
Basiphyllaea
Arpophyllum
Epidendrum
Meiracyllium
Prosthechea
Cattleya
Dilomilis
Restrepia
Pleurothallis
Masdevallia
Anathallis
Polystachya
Angraecum
Aeranthes
Rhipidoglossum
Aerangis
Thrixspermum
Phalaenopsis
Grosourdya
Rhynchostylis
Papilionanthe
Luisia
Vanda
Holcoglossum
Renanthera
Aerides
Arachnis
Pomatocalpa
Gastrochilus
Trichoglottis
Acampe
Vandopsis
Uncifera
Stereochilus
Sarcoglyphis
Smitinandia
Robiquetia
Diploprora
Cleisostoma
Cleisomeria
Schoenorchis
Pelatantheria
Cymbidium
Thecopus
Acriopsis
Dipodium
Galeandra
Catasetum
Dressleria
Mormodes
Cycnoches
Grammatophyllum
Ansellia
Paralophia
Grammangis
Geodorum
Eulophia
Oeceoclades
Eulophiella
Acrolophia
Cyrtopodium
Eriopsis
Zygopetalum
Cryptarrhena
Dichaea
Chaubardia
Rudolfiella
Bifrenaria
Maxillaria
Lycaste
Xylobium
Neomoorea
Peristeria
Lycomormium
Kegeliella
Houlletia
Acineta
Stanhopea
Coryanthes
Rossioglossum
Lockhartia
Cuitlauzina
Fernandezia
Hofmeisterella
Trichoceros
Telipogon
Phymatidium
Zygostates
Chytroglossa
Eloyella
Ornithocephalus
Hintonella
Psychopsis
Trichopilia
Psychopsiella
Grandiphyllum
Trichocentrum
Saundersia
Oncidium
Otoglossum
Gomesa
Rhynchostele
Erycina
Cyrtochiloides
Vitekorchis
Cyrtochilum
Caucaea
Miltonia
Brassia
Cischweinfia
Aspasia
Systeloglossum
Oliveriana
Plectrophora
Zelenkoa
Solenidium
Capanemia
Tolumnia
Notyliopsis
Nohawilliamsia
Leochilus
Pterostemma
Ionopsis
Comparettia
Sutrina
Seegeriella
Rodriguezia
Trizeuxis
Polyotidium
Schunkea
Macradenia
Warmingia
Notylia
Macroclinium
Agrostophyllinae
Coeliinae
CALYPSOEAE
Chysinae
Bletiinae
Laeliinae
EPIDENDREAE
Pleurothallidinae
Polystachyinae
Angraecinae
VANDEAE
Aeridinae
Cymbidiinae*
Cymbidiinae**
Catasetinae
Cymbidiinae***
Eulophiinae
Cyrtopodiinae
Eriopsidinae
Zygopetalinae
Maxillariinae
Coeliopsidinae
Stanhopeinae
CYMBIDIEAE
Oncidiinae
Eocene
Oligocene
Miocene
Pli
Ple
Paleogene
Neogene
Q
50 45 40 35 30 25 20 15 10 5 0
Mya

**Figure S2**

Evolution of growth form and succulent forms (stem and leaves) in epidendroid orchids. (A) Dated phylogenetic tree reconstructed from 203 genera in 14 tribes of epidendroid orchids. Branches with posterior probability < 0.95 are coloured coral-red. Nodes that were constrained in the analysis are indicated by stars (in red when the quartet support for the main topology was < 50% in the coalescent-species tree of Pérez-Escobar *et al*. 2021). (B) and (C) Estimation of ancestral states in the Epidendroideae genus tree with corHMM. As lifestyle and growth form were found to be correlated (see these results in Table 2), these traits were estimated at once using a correlated transition matrix in corHMM. Pie charts at nodes represent ancestral lifestyles and their probabilities. For succulence traits, only significant state changes, i.e. state probability > 0.5 while < 0.5 in the ancestral node, were indicated by coloured triangles (see Supplementary Data Figs. S6 and S8 for full estimations of the ancestral states of stem and leaf succulence respectively). The character states of the present genera are represented on the right side of the tree by coloured boxes. In addition, genera with mycoheterotrophic species are indicated by a mushroom symbol. Cymbidiinae are split into three clades which are indicated by one, two and three stars after the subtribe name.

**Figure S3**

Diagram of transition rates (in events/My) of the correlated lifestyle and growth form computed in corHMM. To note, no genus is recorded to be monopodial and terrestrial in the data, hence the transitions rates reaching the lower bound (1e-9) of the model.

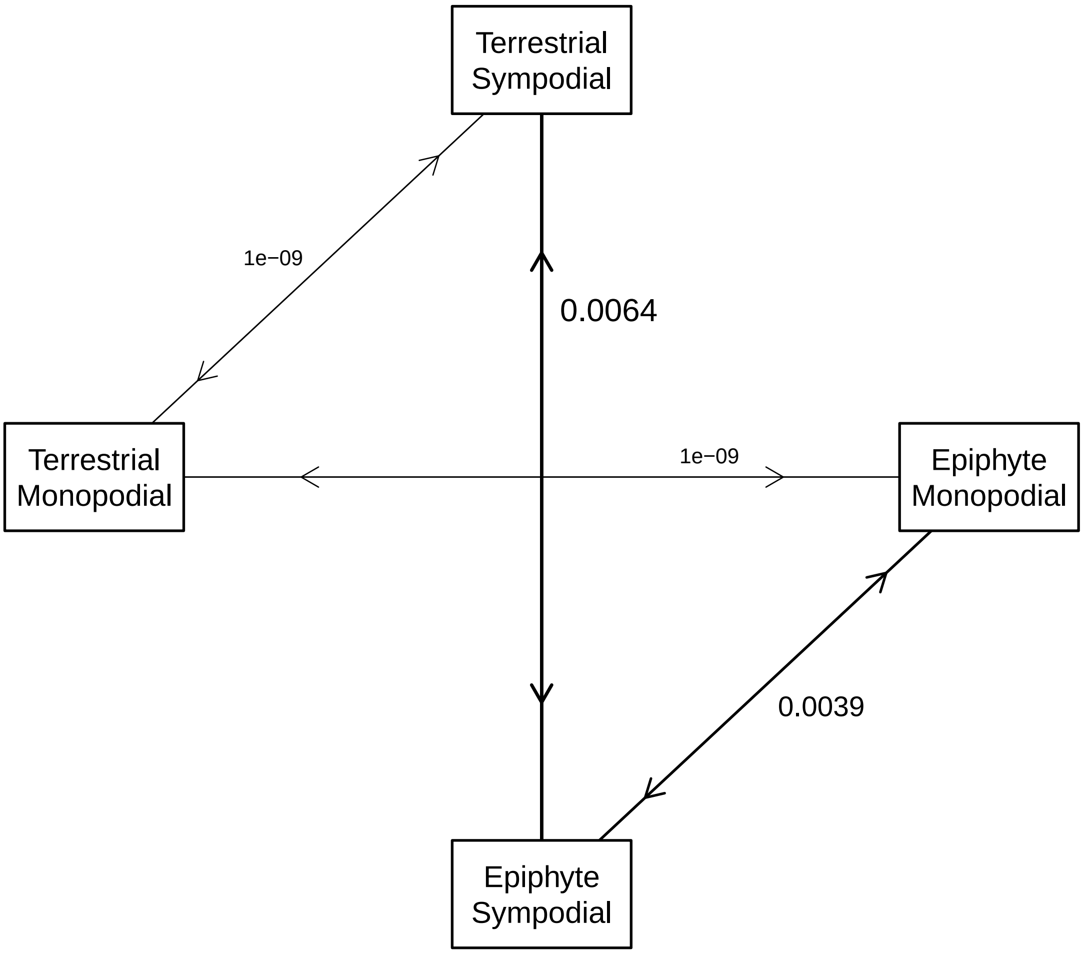

**Figure S4**

Evolution of the position of inflorescence in epidendroid orchids. (A) Dated phylogenetic tree reconstructed from 203 genera in 14 tribes of epidendroid orchids. Branches with posterior probability < 0.95 are coloured coral-red. Nodes that were constrained in the analysis are indicated by stars (in red when the quartet support for the main topology was < 50% in the coalescent-species tree of Pérez-Escobar *et al.* 2021). (B) and (C) Estimation of ancestral states in the Epidendroideae genus tree with corHMM. Pie charts at nodes represent ancestral lifestyles and their probabilities. The character states of the present genera are represented on the right side of the tree by coloured boxes. In addition, genera with mycoheterotrophic species are indicated by a mushroom symbol. Cymbidiinae are split into three clades which are indicated by one, two and three stars after the subtribe name.

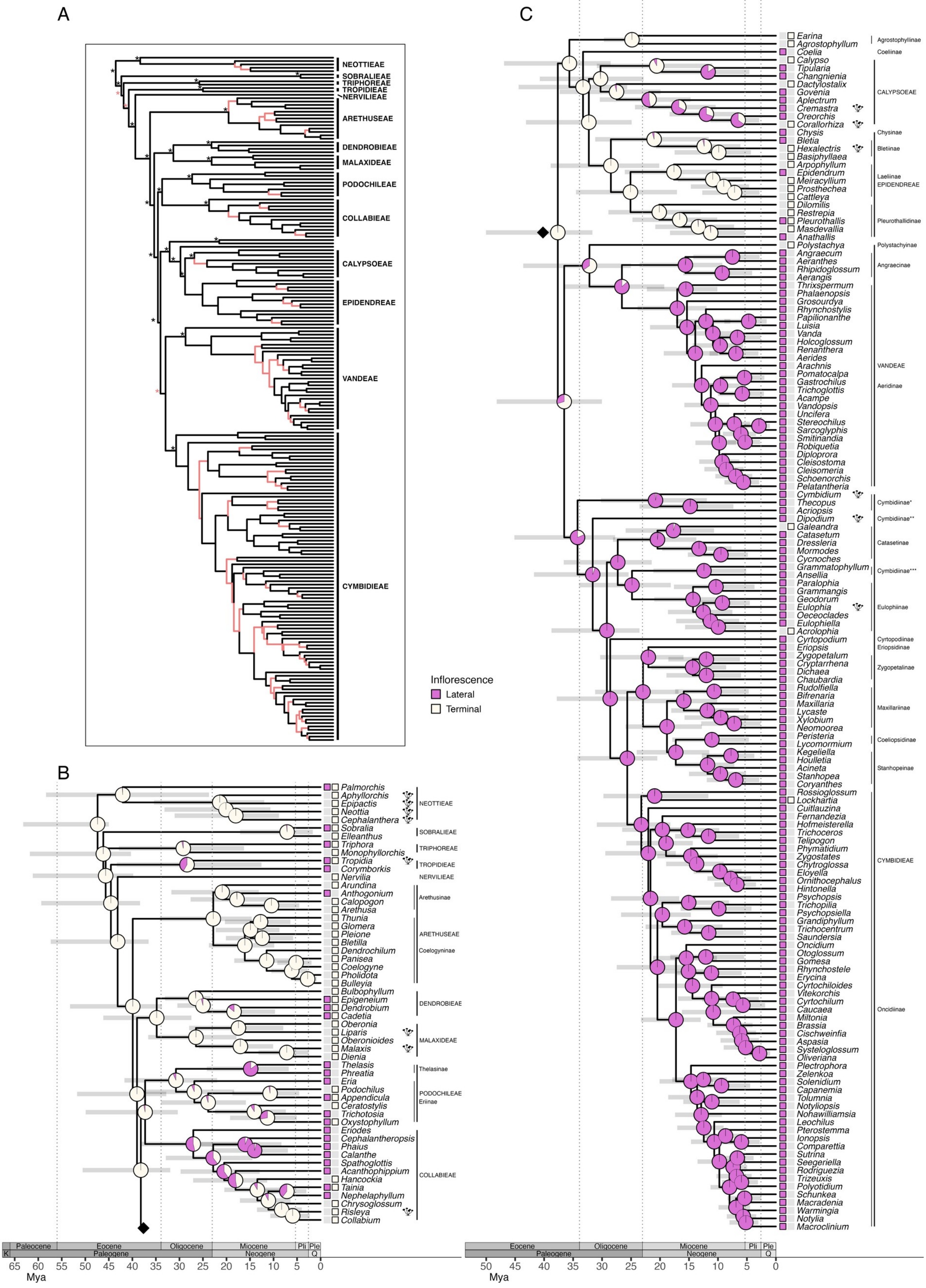
A
NEOTTIEAE
SOBRALIEAE
TRIPHOREAE
TROPIDIEAE
NERVILIEAE
ARETHUSEAE
DENDROBIEAE
MALAXIDEAE
PODOCHILEAE
COLLABIEAE
CALYPSOEAE
EPIDENDREAE
VANDEAE
CYMBIDIEAE
Inflorescence
Lateral
Terminal
B
Palmorchis
Aphyllorchis
Epipactis
Neottia
Cephalanthera
NEOTTIEAE
Sobralia
Elleanthus
SOBRALIEAE
Triphora
Monophyllorchis
TRIPHOREAE
Tropidia
Corymborkis
TROPIDIEAE
Nervilia
NERVILIEAE
Arundina
Anthogonium
Calopogon
Arethusa
Arethusinae
Thunia
Glomera
Pleione
Bletilla
Dendrochilum
Panisea
Coelogyne
Pholidota
Bulleyia
ARETHUSEAE
Coelogyninae
Bulbophyllum
Epigeneium
Dendrobium
Cadetia
DENDROBIEAE
Oberonia
Liparis
Oberonioides
Malaxis
Dienia
MALAXIDEAE
Thelasis
Phreatia
Thelasinae
Eria
Podochilus
Appendicula
Ceratostylis
Trichotosia
Oxystophyllum
PODOCHILEAE
Eriinae
Eriodes
Cephalantheropsis
Phaius
Calanthe
Spathoglottis
Acanthophippium
Hancockia
Tainia
Nephelaphyllum
Chrysoglossum
Risleya
Collabium
COLLABIEAE
K
Paleocene
Eocene
Paleogene
Oligocene
Miocene
Neogene
Pli
Ple
Q
65 60 55 50 45 40 35 30 25 20 15 10 5 0
Mya
C
Earina
Agrostophyllum
Agrostophyllinae
Coelia
Coeliinae
Calypso
Tipularia
Changnienia
Dactylostalix
Govenia
Aplectrum
Cremastra
Oreorchis
Corallorhiza
CALYPSOEAE
Chysis
Chysinae
Bletia
Hexalectris
Basiphyllaea
Bletiinae
Arpophyllum
Epidendrum
Meiracyllium
Prosthechea
Cattleya
Laeliinae
EPIDENDREAE
Dilomilis
Restrepia
Pleurothallis
Masdevallia
Anathallis
Pleurothallidinae
Polystachya
Polystachyinae
Angraecum
Aeranthes
Rhipidoglossum
Aerangis
Angraecinae
Thrixspermum
Phalaenopsis
Grosourdya
Rhynchostylis
Papilionanthe
Luisia
Vanda
Holcoglossum
Renanthera
Aerides
Arachnis
Pomatocalpa
Gastrochilus
Trichoglottis
Acampe
Vandopsis
Uncifera
Stereochilus
Sarcoglyphis
Smitinandia
Robiquetia
Diploprora
Cleisostoma
Cleisomeria
Schoenorchis
Pelatantheria
VANDEAE
Aeridinae
Cymbidium
Thecopus
Acriopsis
Dipodium
Cymbidiinae*
Cymbidiinae**
Galeandra
Catasetum
Dressleria
Mormodes
Cycnoches
Catasetinae
Grammatophyllum
Ansellia
Cymbidiinae***
Paralophia
Grammangis
Geodorum
Eulophia
Oeceoclades
Eulophiella
Acrolophia
Eulophiinae
Cyrtopodium
Cyrtopodiinae
Eriopsis
Eriopsidinae
Zygopetalum
Cryptarrhena
Dichaea
Chaubardia
Zygopetalinae
Rudolfiella
Bifrenaria
Maxillaria
Lycaste
Xylobium
Neomoorea
Maxillariinae
Peristeria
Lycomormium
Coeliopsidinae
Kegeliella
Houlletia
Acineta
Stanhopea
Coryanthes
Stanhopeinae
Rossioglossum
Lockhartia
Cuitlauzina
Fernandezia
Hofmeisterella
Trichoceros
Telipogon
Phymatidium
Zygostates
Chytroglossa
Eloyella
Ornithocephalus
Hintonella
Psychopsis
Trichopilia
Psychopsiella
Grandiphyllum
Trichocentrum
Saundersia
Oncidium
Otoglossum
Gomesa
Rhynchostele
Erycina
Cyrtochiloides
Vitekorchis
Cyrtochilum
Caucaea
Miltonia
Brassia
Cischweinfia
Aspasia
Systeloglossum
Oliveriana
Plectrophora
Zelenkoa
Solenidium
Capanemia
Tolumnia
Notylopsis
Nohawilliamsia
Leochilus
Pterostemma
Ionopsis
Comparettia
Sutrina
Seegeriella
Rodriguezia
Trizeuxis
Polyotidium
Schunkea
Macradenia
Warmingia
Notylia
Macroclinium
CYMBIDIEAE
Oncidiinae
Eocene
Paleogene
Oligocene
Miocene
Neogene
Pli
Ple
Q
50 45 40 35 30 25 20 15 10 5 0
Mya

**Figure S5**

Evolution of the presence/absence of succulent stems in epidendroid orchids. (A) Dated phylogenetic tree reconstructed from 203 genera in 14 tribes of epidendroid orchids. Branches with posterior probability < 0.95 are coloured coral-red. Nodes that were constrained in the analysis are indicated by stars (in red when the quartet support for the main topology was < 50% in the coalescent-species tree of Pérez-Escobar *et al*. 2021). (B) and (C) Estimation of ancestral states in the Epidendroideae genus tree with corHMM. Pie charts at nodes represent ancestral lifestyles and their probabilities. The character states of the present genera are represented on the right side of the tree by coloured boxes. In addition, genera with mycoheterotrophic species are indicated by a mushroom symbol. Cymbidiinae are split into three clades which are indicated by one, two and three stars after the subtribe name.

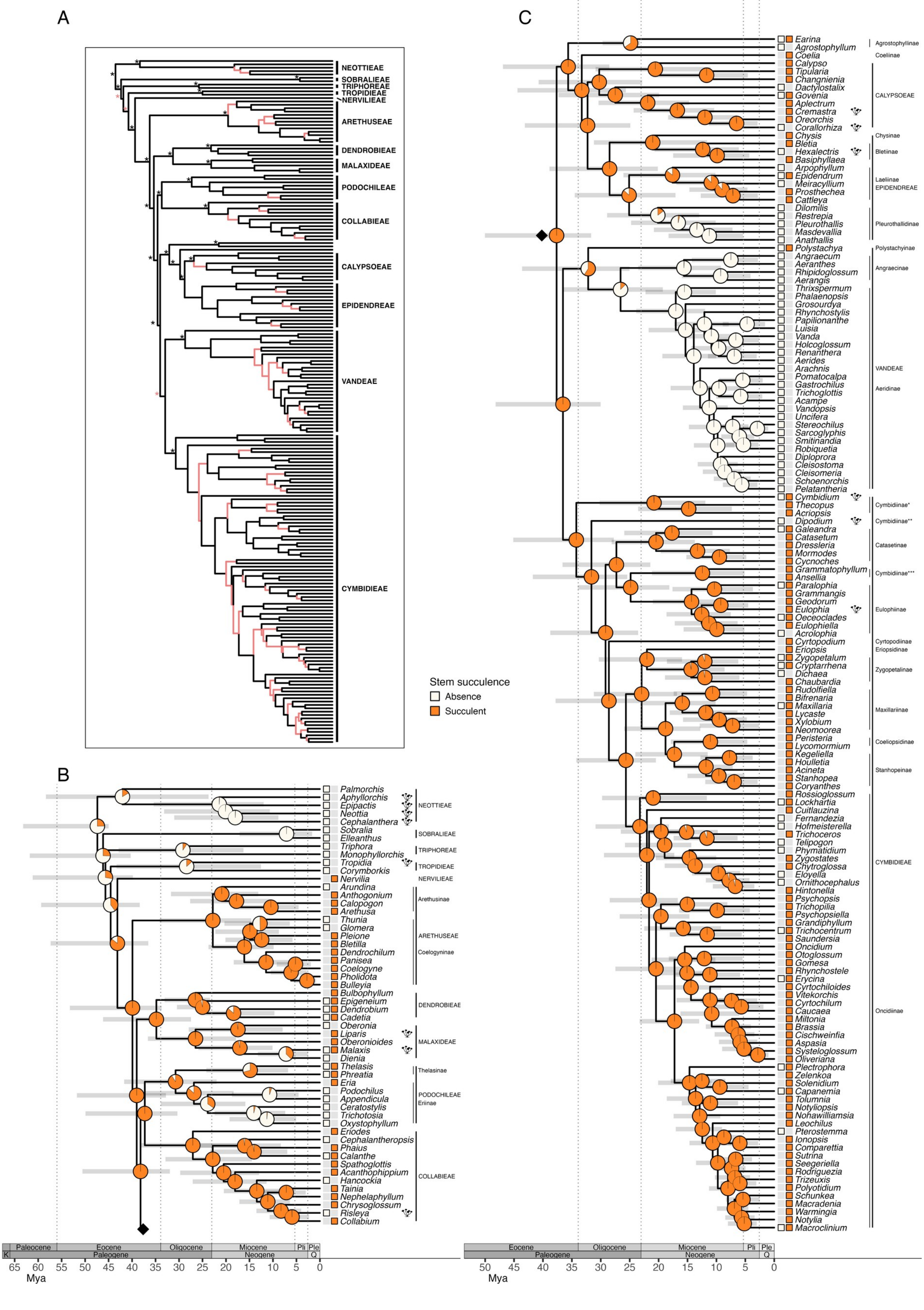
A
NEOTTIEAE
SOBRALIEAE
TRIPHOREAE
TROPIDIEAE
NERVILIEAE
ARETHUSEAE
DENDROBIEAE
MALAXIDEAE
PODOCHILEAE
COLLABIEAE
CALYPSOEAE
EPIDENDREAE
VANDEAE
CYMBIDIEAE
Stem succulence
Absence
Succulent
B
Palmorchis
Aphyllorchis
Epipactis
Neottia
Cephalanthera
Sobralia
Elleanthus
Triphora
Monophyllorchis
Tropidia
Corymborkis
Nervilia
Arundina
Anthogonium
Calopogon
Arethusa
Thunia
Glomera
Pleione
Bletilla
Dendrochilum
Panisea
Coelogyne
Pholidota
Bulleyia
Bulbophyllum
Epigeneium
Dendrobium
Cadetia
Oberonia
Liparis
Oberonioides
Malaxis
Dienia
Thelasis
Phreatia
Eria
Podochilus
Appendicula
Ceratostylis
Trichotosia
Oxystophyllum
Eriodes
Cephalantheropsis
Phaius
Calanthe
Spathoglottis
Acanthophippium
Hancockia
Tainia
Nephelaphyllum
Chrysoglossum
Risleya
Collabium
NEOTTIEAE
SOBRALIEAE
TRIPHOREAE
TROPIDIEAE
NERVILIEAE
Arethusinae
ARETHUSEAE
Coelogyninae
DENDROBIEAE
MALAXIDEAE
Thelasinae
PODOCHILEAE
Eriinae
COLLABIEAE
Paleocene
Eocene
Oligocene
Miocene
Pli
Ple
K
Paleogene
Neogene
Q
65 60 55 50 45 40 35 30 25 20 15 10 5 0
Mya
C
Earina
Agrostophyllum
Coelia
Calypso
Tipularia
Changnienia
Dactylostalix
Govenia
Aplectrum
Cremastra
Oreorchis
Corallorhiza
Chysis
Bletia
Hexalectris
Basiphyllaea
Arpophyllum
Epidendrum
Meiracyllium
Prosthechea
Cattleya
Dilomilis
Restrepia
Pleurothallis
Masdevallia
Anathallis
Polystachya
Angraecum
Aeranthes
Rhipidoglossum
Aerangis
Thrixspermum
Phalaenopsis
Grosourdya
Rhynchostylis
Papilionanthe
Luisia
Vanda
Holcoglossum
Renanthera
Aerides
Arachnis
Pomatocalpa
Gastrochilus
Trichoglottis
Acampe
Vandopsis
Uncifera
Stereochilus
Sarcoglyphis
Smitinandia
Robiquetia
Diploprora
Cleisostoma
Cleisomeria
Schoenorchis
Pelatantheria
Cymbidium
Thecopus
Acriopsis
Dipodium
Galeandra
Catasetum
Dressleria
Mormodes
Cycnoches
Grammatophyllum
Ansellia
Paralophia
Grammangis
Geodorum
Eulophia
Oeceoclades
Eulophiella
Acrolophia
Cyrtopodium
Eriopsis
Zygopetalum
Cryptarrhena
Dichaea
Chaubardia
Rudolfiella
Bifrenaria
Maxillaria
Lycaste
Xylobium
Neomoorea
Peristeria
Lycomormium
Kegeliella
Houlletia
Acineta
Stanhopea
Coryanthes
Rossioglossum
Lockhartia
Cuitlauzina
Fernandezia
Hofmeisterella
Trichoceros
Telipogon
Phymatidium
Zygostates
Chytroglossa
Eloyella
Ornithocephalus
Hintonella
Psychopsis
Trichopilia
Psychopsiella
Grandiphyllum
Trichocentrum
Saundersia
Oncidium
Otoglossum
Gomesa
Rhynchostele
Erycina
Cyrtochiloides
Vitekorchis
Cyrtochilum
Caucaea
Miltonia
Brassia
Cischweinfia
Aspasia
Systeloglossum
Oliveriana
Plectrophora
Zelenkoa
Solenidium
Capanemia
Tolumnia
Notyliopsis
Nohawilliamsia
Leochilus
Pterostemma
Ionopsis
Comparettia
Sutrina
Seegeriella
Rodriguezia
Trizeuxis
Polyotidium
Schunkea
Macradenia
Warmingia
Notylia
Macroclinium
Agrostophyllinae
Coeliinae
CALYPSOEAE
Chysinae
Bletiinae
Laeliinae
EPIDENDREAE
Pleurothallidinae
Polystachyinae
Angraecinae
VANDEAE
Aeridinae
Cymbidiinae*
Cymbidiinae**
Catasetinae
Cymbidiinae***
Eulophiinae
Cyrtopodiinae
Eriopsidinae
Zygopetalinae
Maxillariinae
Coeliopsidinae
Stanhopeinae
CYMBIDIEAE
Oncidiinae
Eocene
Oligocene
Miocene
Pli
Ple
Paleogene
Neogene
Q
50 45 40 35 30 25 20 15 10 5 0
Mya

**Figure S6**

Evolution of succulent stems in epidendroid orchids. (A) Dated phylogenetic tree reconstructed from 203 genera in 14 tribes of epidendroid orchids. Branches with posterior probability < 0.95 are coloured coral-red. Nodes that were constrained in the analysis are indicated by stars (in red when the quartet support for the main topology was < 50% in the coalescent-species tree of Pérez-Escobar *et al.* 2021). (B) and (C) Estimation of ancestral states in the Epidendroideae genus tree with corHMM. Pie charts at nodes represent ancestral lifestyles and their probabilities. The character states of the present genera are represented on the right side of the tree by coloured boxes. In addition, genera with mycoheterotrophic species are indicated by a mushroom symbol. Cymbidiinae are split into three clades which are indicated by one, two and three stars after the subtribe name.

**Figure S7**

Diagram of transition rates (in events/My) of the stem succulence computed in corHMM.

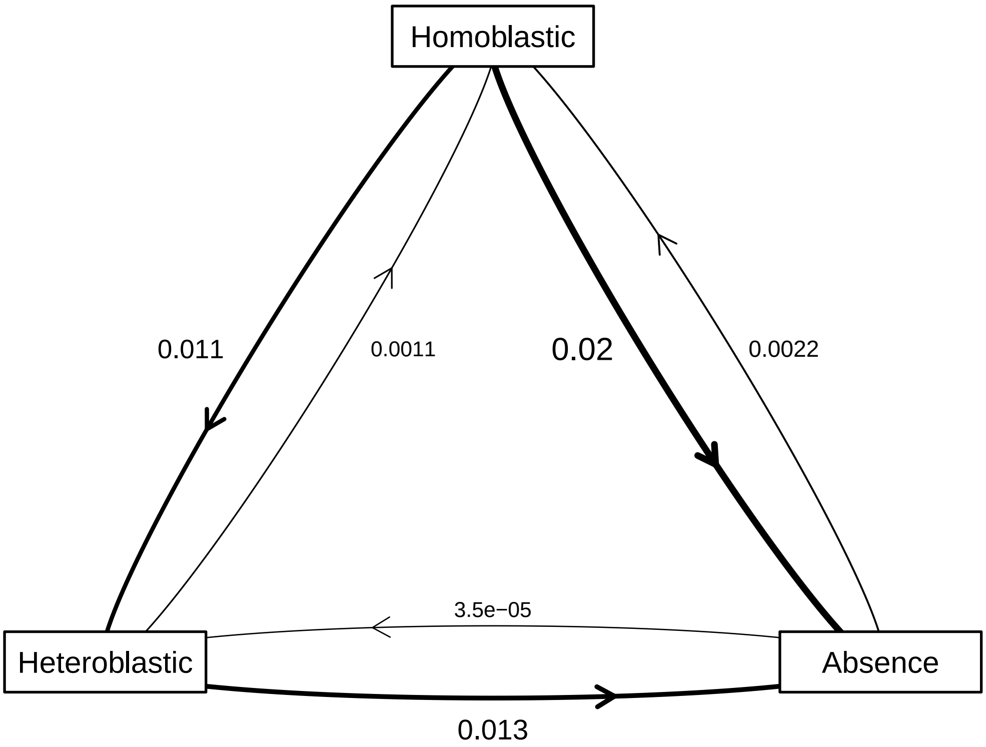

**Figure S8**

Evolution of lifestyle and succulent leaves in epidendroid orchids. (A) Dated phylogenetic tree reconstructed from 203 genera in 14 tribes of epidendroid orchids. Branches with posterior probability < 0.95 are coloured coral-red. Nodes that were constrained in the analysis are indicated by stars (in red when the quartet support for the main topology was < 50% in the coalescent-species tree of Pérez-Escobar *et al.* 2021). (B) and (C) Estimation of ancestral states in the Epidendroideae genus tree with corHMM. As lifestyle and leaf succulence were found to be correlated (see these results in Table 2), these traits were estimated at once using a correlated transition matrix in corHMM. Pie charts at nodes represent ancestral lifestyles and their probabilities. For lifestyle, only significant state changes, i.e. state probability > 0.5 while < 0.5 in the ancestral node, were indicated by coloured triangles (see Fig. 2 for full estimations of the ancestral states of lifestyle). The character states of the present genera are represented on the right side of the tree by coloured boxes. In addition, genera with mycoheterotrophic species are indicated by a mushroom symbol. Cymbidiinae are split into three clades which are indicated by one, two and three stars after the subtribe name.

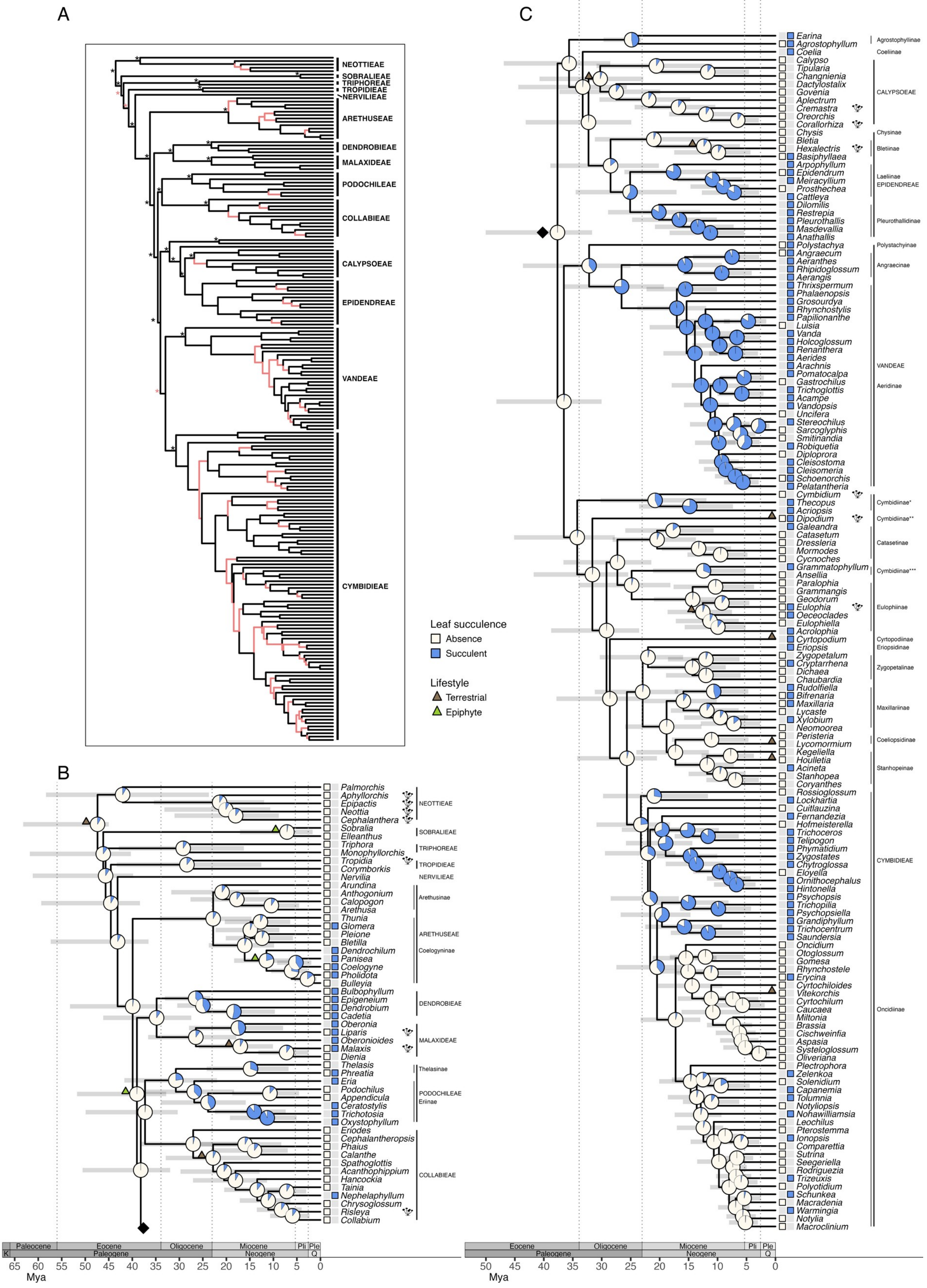
A
NEOTTIEAE
SOBRALIEAE
TRIPHOREAE
TROPIDIEAE
NERVILIEAE
ARETHUSEAE
DENDROBIEAE
MALAXIDEAE
PODOCHILEAE
COLLABIEAE
CALYPSOEAE
EPIDENDREAE
VANDEAE
CYMBIDIEAE
Leaf succulence
Absence
Succulent
Lifestyle
Terrestrial
Epiphyte
B
Palmorchis
Aphyllorchis
Epipactis
Neottia
Cephalanthera
NEOTTIEAE
Sobralia
Elleanthus
SOBRALIEAE
Triphora
Monophyllorchis
TRIPHOREAE
Tropidia
Corymborkis
TROPIDIEAE
Nervilia
NERVILIEAE
Arundina
Anthogonium
Calopogon
Arethusa
Arethusinae
Thunia
Glomera
Pleione
Bletilla
Dendrochilum
Panisea
Coelogyne
Pholidota
Bulleyia
ARETHUSEAE
Coelogyninae
Bulbophyllum
Epigeneium
Dendrobium
Cadetia
DENDROBIEAE
Oberonia
Liparis
Oberonioides
Malaxis
Dienia
MALAXIDEAE
Thelasis
Phreatia
Thelasinae
Eria
Podochilus
Appendicula
Ceratostylis
Trichotosia
Oxystophyllum
PODOCHILEAE
Eriinae
Eriodes
Cephalantheropsis
Phaius
Calanthe
Spathoglottis
Acanthophippium
Hancockia
Tainia
Nephelaphyllum
Chrysoglossum
Risleya
Collabium
COLLABIEAE
Paleocene
Eocene
Oligocene
Miocene
Pli
Ple
K
Paleogene
Neogene
Q
65 60 55 50 45 40 35 30 25 20 15 10 5 0
Mya
C
Earina
Agrostophyllum
Agrostophyllinae
Coelia
Coeliinae
Calypso
Tipularia
Changnienia
Dactylostalix
Govenia
Aplectrum
Cremastra
Oreorchis
Corallorhiza
CALYPSOEAE
Chysis
Chysinae
Bletia
Hexalectris
Basiphyllaea
Bletiinae
Arpophyllum
Epidendrum
Meiracyllium
Prosthechea
Cattleya
Laeliinae
EPIDENDREAE
Dilomilis
Restrepia
Pleurothallis
Masdevallia
Anathallis
Pleurothallidinae
Polystachya
Polystachyinae
Angraecum
Aeranthes
Rhipidoglossum
Aerangis
Angraecinae
Thrixspermum
Phalaenopsis
Grosourdya
Rhynchostylis
Papilionanthe
Luisia
Vanda
Holcoglossum
Renanthera
Aerides
Arachnis
Pomatocalpa
Gastrochilus
Trichoglottis
Acampe
Vandopsis
Uncifera
Stereochilus
Sarcoglyphis
Smitinandia
Robiquetia
Diploprora
Cleisostoma
Cleisomeria
Schoenorchis
Pelatantheria
VANDEAE
Aeridinae
Cymbidium
Thecopus
Acriopsis
Dipodium
Galeandra
Cymbidiinae*
Cymbidiinae**
Catasetum
Dressleria
Mormodes
Cycnoches
Catasetinae
Grammatophyllum
Ansellia
Cymbidiinae***
Paralophia
Grammangis
Geodorum
Eulophia
Oeceoclades
Eulophiella
Eulophiinae
Acrolophia
Cyrtopodium
Eriopsis
Cyrtopodiinae
Eriopsidinae
Zygopetalum
Cryptarrhena
Dichaea
Chaubardia
Zygopetalinae
Rudolfiella
Bifrenaria
Maxillaria
Lycaste
Xylobium
Neomoorea
Maxillariinae
Peristeria
Lycomormium
Coeliopsidinae
Kegeliella
Houlletia
Acineta
Stanhopea
Coryanthes
Stanhopeinae
Rossioglossum
Lockhartia
Cuitlauzina
Fernandezia
Hofmeisterella
Trichoceros
Telipogon
Phymatidium
Zygostates
Chytroglossa
Eloyella
Ornithocephalus
Hintonella
Psychopsis
Trichopilia
Psychopsiella
Grandiphyllum
Trichocentrum
Saundersia
Oncidium
Otoglossum
Gomesa
Rhynchostele
Erycina
Cyrtochiloides
Vitekorchis
Cyrtochilum
Caucaea
Miltonia
Brassia
Cischweinfia
Aspasia
Systeloglossum
Oliveriana
Plectrophora
Zelenkoa
Solenidium
Capanemia
Tolumnia
Notyliopsis
Nohawilliamsia
Leochilus
Pterostemma
Ionopsis
Comparettia
Sutrina
Seegeriella
Rodriguezia
Trizeuxis
Polyotidium
Schunkea
Macradenia
Warmingia
Notylia
Macroclinium
CYMBIDIEAE
Oncidiinae
Eocene
Oligocene
Miocene
Pli
Ple
Paleogene
Neogene
Q
50 45 40 35 30 25 20 15 10 5 0
Mya

**Figure S9**

Evolution of the mean number of velamen layers in epidendroid orchids. (A) Dated phylogenetic tree reconstructed from 203 genera in 14 tribes of epidendroid orchids. Branches with posterior probability < 0.95 are coloured coral-red. Nodes that were constrained in the analysis are indicated by stars (in red when the quartet support for the main topology was < 50% in the coalescent-species tree of Pérez-Escobar *et al*. 2021). (B) and (C) Estimation of ancestral states in the Epidendroideae genus tree with anc.ML. Missing data for the number of velamen layers are indicated by stars at tips before the genus names. Genera missing data on the number of velamen layers inheriting the ancestral state without change, hence some genera missing data may actually have a lower or a higher number of velamen layers. In addition, genera with mycoheterotrophic species are indicated by a mushroom symbol. Cymbidiinae are split into three clades which are indicated by one, two and three stars after the subtribe name.

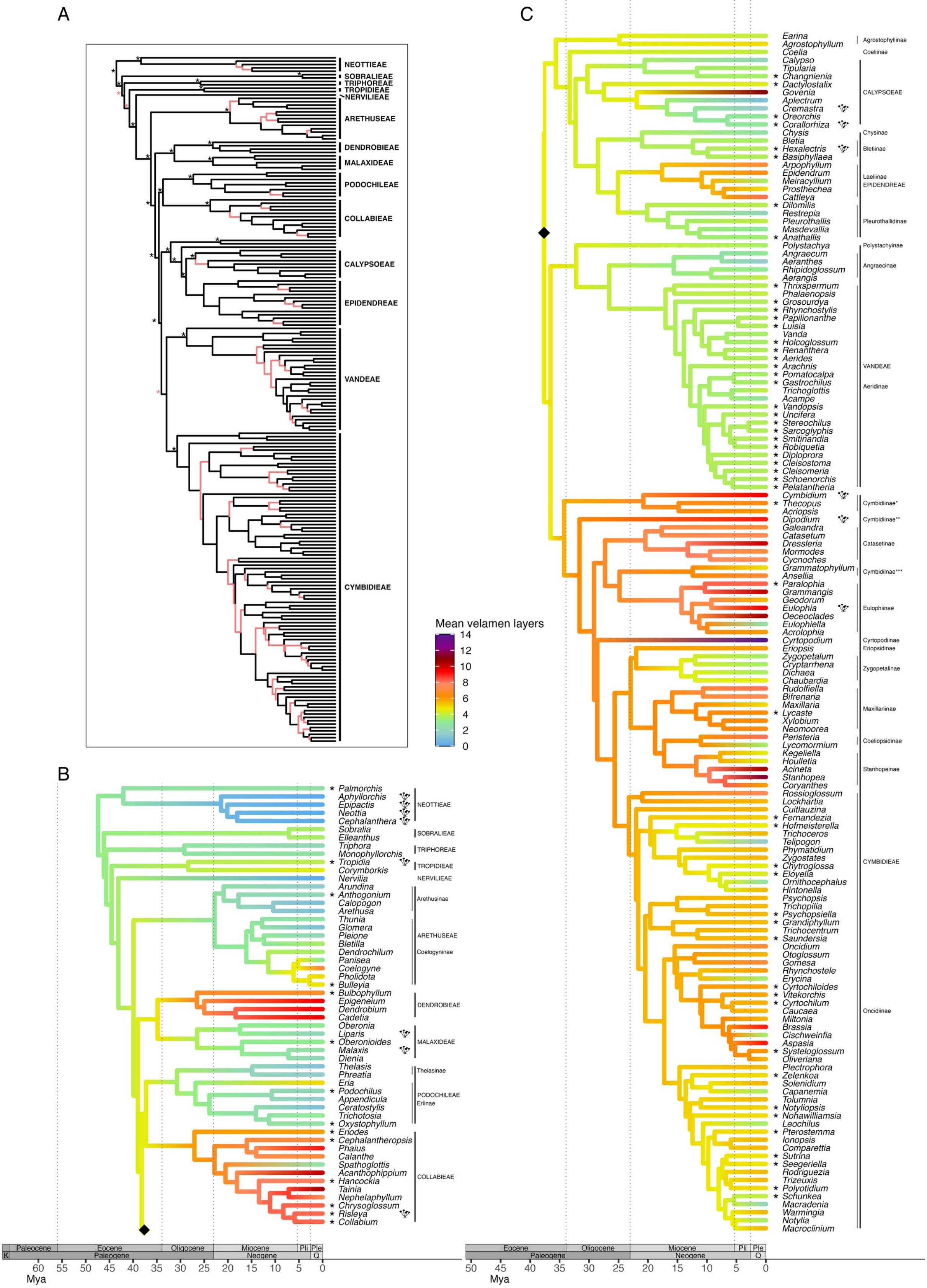

A
NEOTTIEAE
SOBRALIEAE
TRIPHOREAE
TROPIDIEAE
NERVILIEAE
ARETHUSEAE
DENDROBIEAE
MALAXIDEAE
PODOCHILEAE
COLLABIEAE
CALYPSOEAE
EPIDENDREAE
VANDEAE
CYMBIDIEAE
Mean velamen layers
14
12
10
8
6
4
2
0
B
Palmorchis
Aphyllorchis
Epipactis
Neottia
Cephalanthera
NEOTTIEAE
Sobralia
Elleanthus
SOBRALIEAE
Triphora
Monophyllorchis
TRIPHOREAE
Tropidia
Corymborkis
TROPIDIEAE
Nervilia
NERVILIEAE
Arundina
Anthogonium
Calopogon
Arethusa
Arethusinae
Thunia
Glomera
Pleione
Bletilla
Dendrochilum
Panisea
Coelogyne
Pholidota
Bulleyia
ARETHUSEAE
Coelogyninae
Bulbophyllum
Epigeneium
Dendrobium
Cadetia
DENDROBIEAE
Oberonia
Liparis
Oberonioides
Malaxis
Dienia
MALAXIDEAE
Thelasis
Phreatia
Thelasinae
Eria
Podochilus
Appendicula
Ceratostylis
Trichotosia
Oxystophyllum
PODOCHILEAE
Eriinae
Eriodes
Cephalantheropsis
Phaius
Calanthe
Spathoglottis
Acanthophippium
Hancockia
Tainia
Nephelaphyllum
Chrysoglossum
Risleya
Collabium
COLLABIEAE
Paleocene
Eocene
Oligocene
Miocene
Pli
Ple
K
Paleogene
Neogene
Q
60
55
50
45
40
35
30
25
20
15
10
5
0
Mya
C
Earina
Agrostophyllum
Agrostophyllinae
Coelia
Coeliinae
Calypso
Tipularia
Changnienia
Dactylostalix
Govenia
Aplectrum
Cremastra
Oreorchis
Corallorhiza
CALYPSOEAE
Chysis
Chysinae
Bletia
Hexalectris
Basiphyllaea
Bletiinae
Arpophyllum
Epidendrum
Meiracyllium
Prosthechea
Cattleya
Laeliinae
EPIDENDREAE
Dilomilis
Restrepia
Pleurothallis
Masdevallia
Anathallis
Pleurothallidinae
Polystachya
Polystachyinae
Angraecum
Aeranthes
Rhipidoglossum
Aerangis
Angraecinae
Thrixspermum
Phalaenopsis
Grosourdya
Rhynchostylis
Papilionanthe
Luisia
Vanda
Holcoglossum
Renanthera
Aerides
Arachnis
Pomatocalpa
Gastrochilus
Trichoglottis
Acampe
Vandopsis
Uncifera
Stereochilus
Sarcoglyphis
Smitinandia
Robiquetia
Diploprora
Cleisostoma
Cleisomeria
Schoenorchis
Pelatantheria
VANDEAE
Aeridinae
Cymbidium
Thecopus
Acriopsis
Cymbidiinae*
Dipodium
Cymbidiinae**
Galeandra
Catasetum
Dressleria
Mormodes
Cycnoches
Catasetinae
Grammatophyllum
Ansellia
Cymbidiinae***
Paralophia
Grammangis
Geodorum
Eulophia
Oeceoclades
Eulophiella
Acrolophia
Eulophiinae
Cyrtopodium
Cyrtopodiinae
Eriopsis
Eriopsidinae
Zygopetalum
Cryptarrhena
Dichaea
Chaubardia
Zygopetalinae
Rudolfiella
Bifrenaria
Maxillaria
Lycaste
Xylobium
Neomoorea
Maxillariinae
Peristeria
Lycomormium
Coeliopsidinae
Kegeliella
Houlletia
Acineta
Stanhopea
Coryanthes
Stanhopeinae
Rossioglossum
Lockhartia
Cuitlauzina
Fernandezia
Hofmeisterella
Trichoceros
Telipogon
Phymatidium
Zygostates
Chytroglossa
Eloyella
Ornithocephalus
Hintonella
Psychopsis
Trichopilia
Psychopsiella
Grandiphyllum
Trichocentrum
Saundersia
Oncidium
Otoglossum
Gomesa
Rhynchostele
Erycina
Cyrtochiloides
Vitekorchis
Cyrtochilum
Caucaea
Miltonia
Brassia
Cischweinfia
Aspasia
Systeloglossum
Oliveriana
Plectrophora
Zelenkoa
Solenidium
Capanemia
Tolumnia
Notyliopsis
Nohawilliamsia
Leochilus
Pterostemma
Ionopsis
Comparettia
Sutrina
Seegeriella
Rodriguezia
Trizeuxis
Polyotidium
Schunkea
Macradenia
Warmingia
Notylia
Macroclinium
CYMBIDIEAE
Oncidiinae
Eocene
Oligocene
Miocene
Pli
Ple
Paleogene
Neogene
Q
50
45
40
35
30
25
20
15
10
5
0
Mya

**Figure S10**

Lineage-through-time plot of terrestrial genera of Epidendroideae sampled in the phylogeny (75 genera).

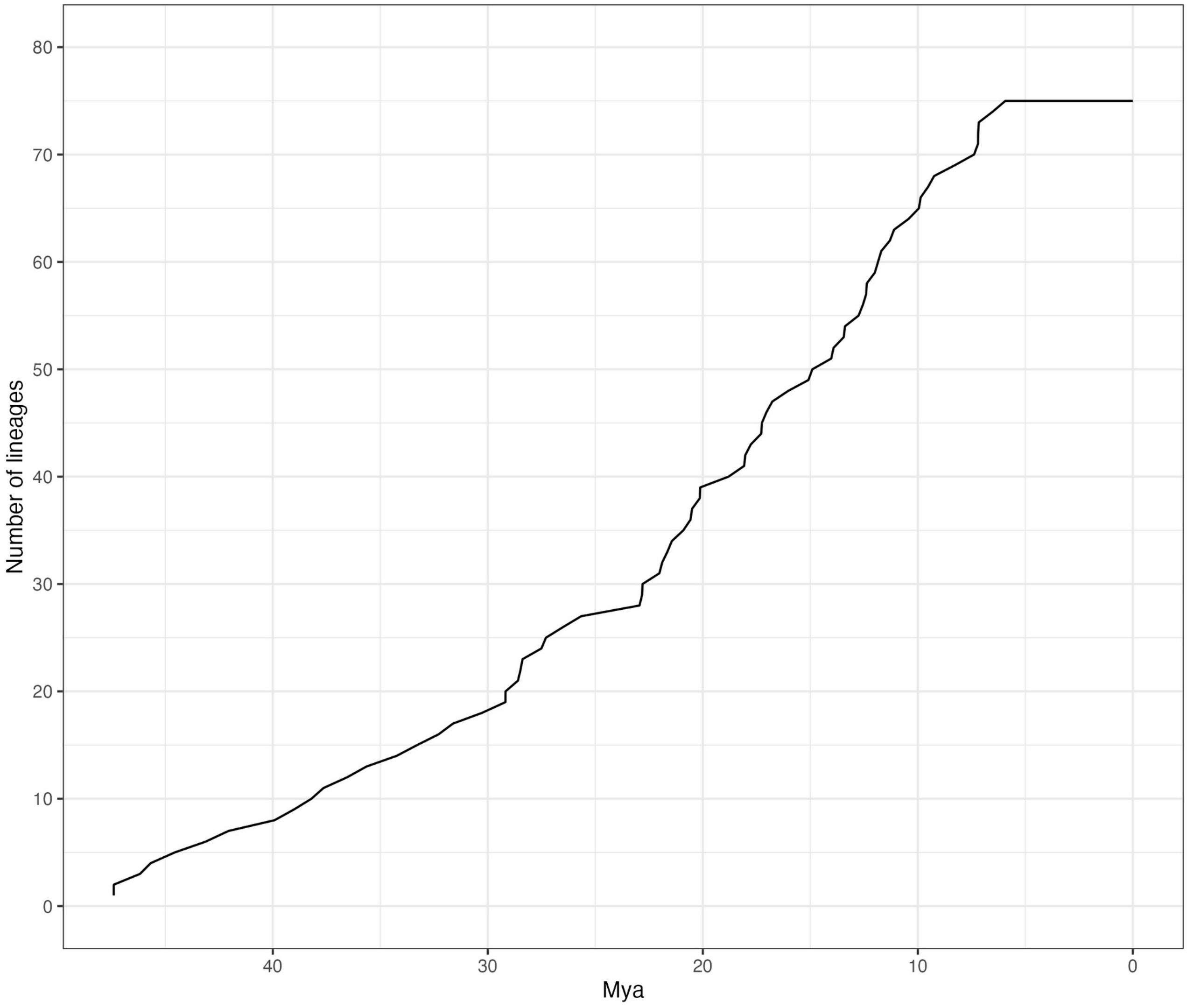

**Figure S11**

Lineage-through-time plot of epiphytic genera of Epidendroideae sampled in the phylogeny (153 genera).

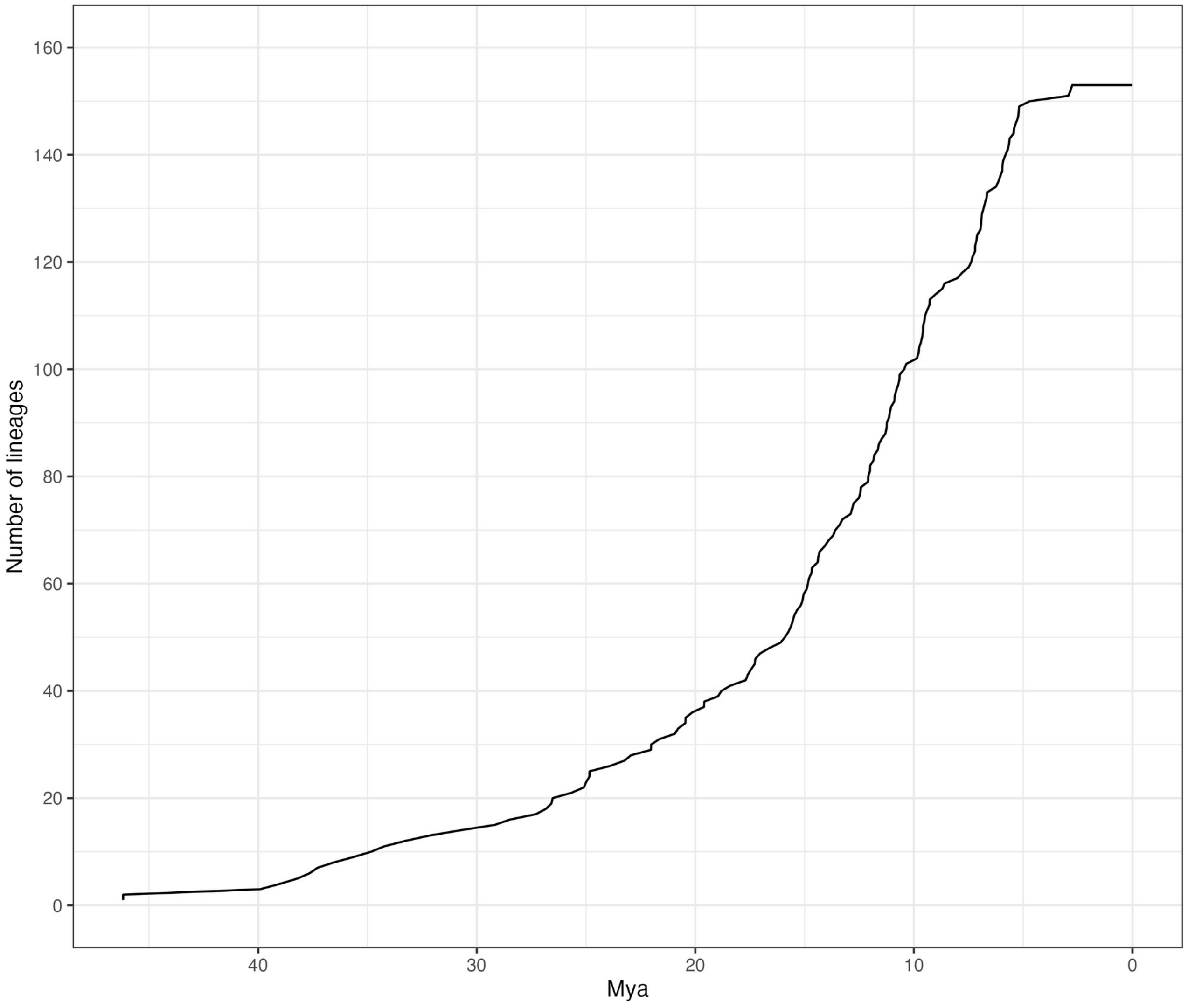
